# Mathematical modeling of plus-strand RNA virus replication to identify broad-spectrum antiviral treatment strategies

**DOI:** 10.1101/2022.07.25.501353

**Authors:** Carolin Zitzmann, Christopher Dächert, Bianca Schmid, Hilde van der Schaar, Martijn van Hemert, Alan S. Perelson, Frank J.M. van Kuppeveld, Ralf Bartenschlager, Marco Binder, Lars Kaderali

## Abstract

Plus-strand RNA viruses are the largest group of viruses. Many are human pathogens that inflict a socio-economic burden. Interestingly, plus-strand RNA viruses share remarkable similarities in their replication. A hallmark of plus-strand RNA viruses is the remodeling of intracellular membranes to establish replication organelles (so-called “replication factories”), which provide a protected environment for the replicase complex, consisting of the viral genome and proteins necessary for viral RNA synthesis. In the current study, we investigate pan-viral similarities and virus-specific differences in the life cycle of this highly relevant group of viruses. We first measured the kinetics of viral RNA, viral protein, and infectious virus particle production of hepatitis C virus (HCV), dengue virus (DENV), and coxsackievirus B3 (CVB3) in the immuno-compromised Huh7 cell line and thus without perturbations by an intrinsic immune response. Based on these measurements, we developed a detailed mathematical model of the replication of HCV, DENV, and CVB3 and show that only small virus-specific changes in the model were necessary to describe the *in vitro* dynamics of the different viruses. Our model correctly predicted virus-specific mechanisms such as host cell translation shut off and different kinetics of replication organelles. Further, our model suggests that the ability to suppress or shut down host cell mRNA translation may be a key factor for *in vitro* replication efficiency which may determine acute self-limited or chronic infection. We further analyzed potential broad-spectrum antiviral treatment options *in silico* and found that targeting viral RNA translation, especially polyprotein cleavage, and viral RNA synthesis may be the most promising drug targets for all plus-strand RNA viruses. Moreover, we found that targeting only the formation of replicase complexes did not stop the viral replication *in vitro* early in infection, while inhibiting intracellular trafficking processes may even lead to amplified viral growth.

**Author summary:** Plus-strand RNA viruses comprise a large group of related and medically relevant viruses. The current global pandemic of COVID-19 caused by the SARS-coronavirus-2 as well as the constant spread of diseases such as dengue and chikungunya fever show the necessity of a comprehensive and precise analysis of plus-strand RNA virus infections. Plus-strand RNA viruses share similarities in their life cycle. To understand their within-host replication strategies, we developed a mathematical model that studies pan-viral similarities and virus-specific differences of three plus-strand RNA viruses, namely hepatitis C, dengue, and coxsackievirus. By fitting our model to *in vitro* data, we found that only small virus-specific variations in the model were required to describe the dynamics of all three viruses. Furthermore, our model predicted that ribosomes involved in viral RNA translation seem to be a key player in plus-strand RNA replication efficiency, which may determine acute or chronic infection outcome. Furthermore, our *in-silico* drug treatment analysis suggests that targeting viral proteases involved in polyprotein cleavage, in combination with viral RNA replication, may represent promising drug targets with broad-spectrum antiviral activity.

## Introduction

Plus-strand RNA viruses are the largest group of human pathogens that cause re-emerging epidemics as seen with dengue, chikungunya and Zika virus, as well as global pandemics of acute and chronic infectious diseases such as hepatitis C and the common cold. The current global SARS-coronavirus-2 (SARS-CoV-2) pandemic shows how our lives can become affected by a rapidly spreading plus-strand RNA virus. As of May 2022, more than 500 million cases of SARS-CoV-2 infections have been reported with over 6 million confirmed deaths [1,2]. While a global pandemic of the current scale clearly causes an exceptional socio-economic burden [3], various other plus-strand RNA viruses cause significant burden as well. For example, in 2013, symptomatic dengue cases in 141 countries caused socio-economic costs of US$ 8.9 billion [4], while the costs of the latest Zika outbreak has been estimated as US$ 7-18 billion in Latin America and the Caribbean from 2015 to 2017 [5]. Furthermore, between 2014 and 2018, the USA spend around US$ 60 billion for hepatitis C medication with around US$ 80,000 per patient [6,7].

Treatment options are limited for the majority of plus-strand RNA viruses. While there are vaccines and vaccine candidates available for few viruses, approved direct acting antivirals are only available against hepatitis C and SARS-CoV-2 [8,9]. Given the high disease burden and socio-economic cost caused by infections with plus-strand RNA viruses, there is an urgent need for broadly acting antiviral drugs. To develop these, it is important to study the life cycles and host restriction and dependency factors in detail, not only at the level of each virus individually, but also across a group of related viruses to gain pan-viral insights. In the current study, we investigated the life cycle of plus-strand RNA viruses. The ultimate goal was to reveal commonly effective antiviral strategies and potential therapeutic target processes in the viral life cycle. To do so, we chose three representatives of plus-strand RNA viruses, hepatitis C, dengue, and coxsackievirus B3 (compare Table 1).

**Table 1:**
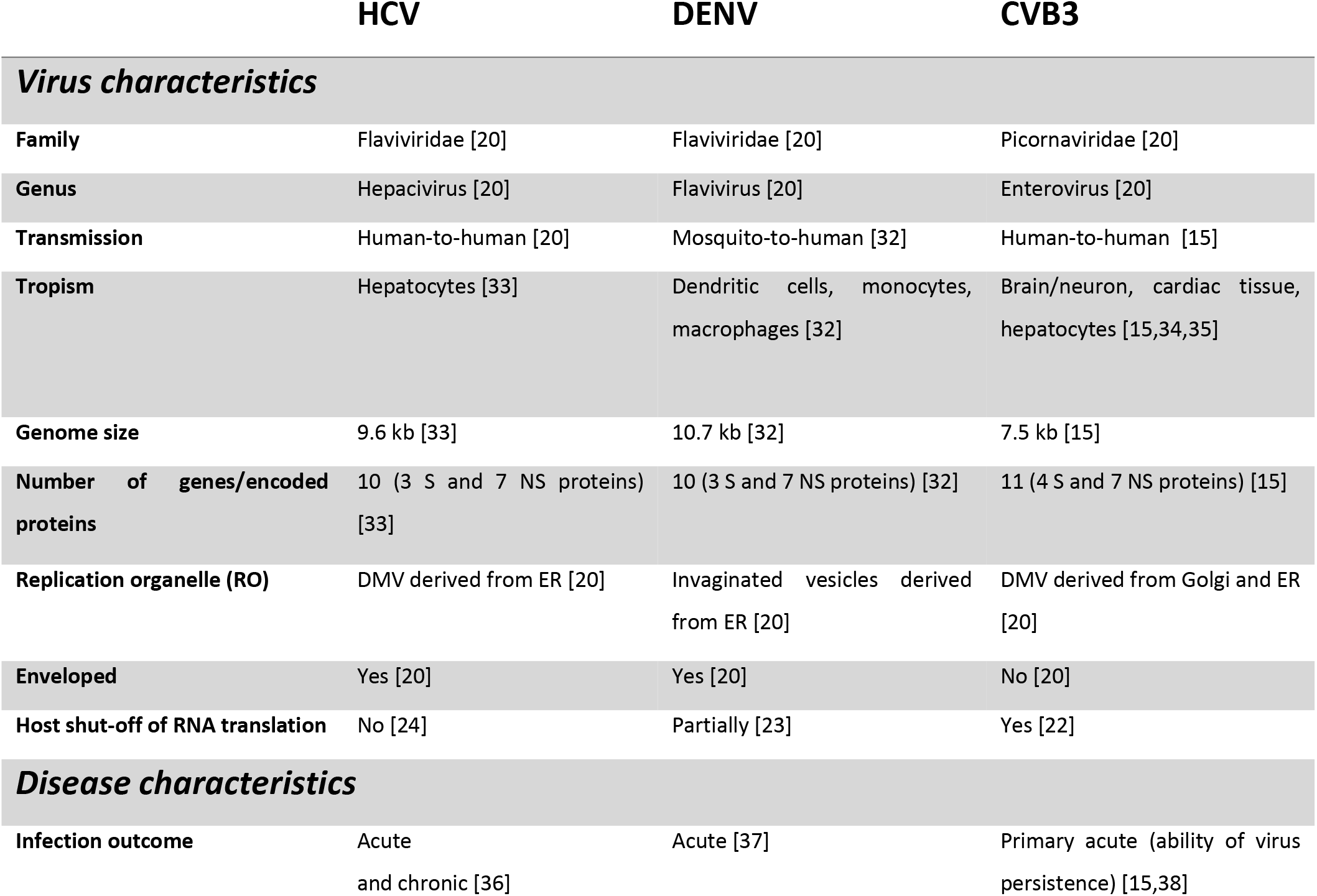
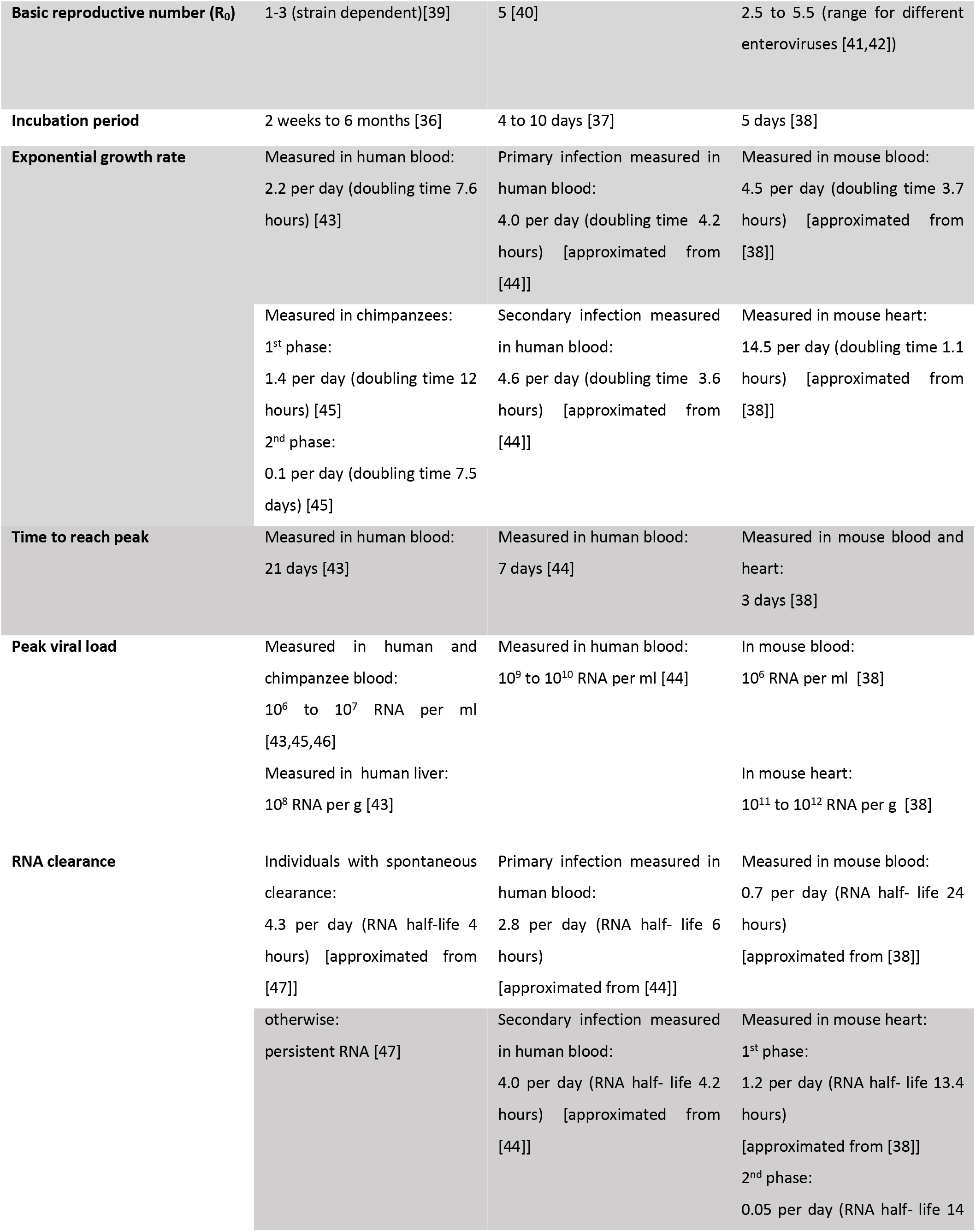
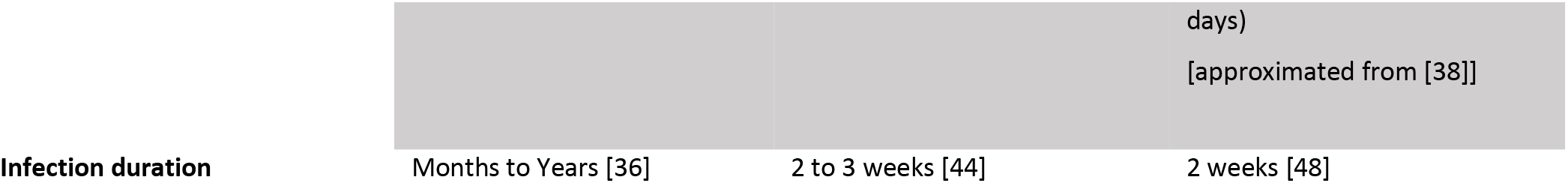
Feature comparison of plus-strand RNA viruses. DMV: double membrane vesicles, ER: endoplasmic reticulum, NS: non-structural, S: structural.

The enveloped blood-borne hepatitis C virus (HCV) is a *Hepacivirus* of the family *Flaviviridae* that causes acute and chronic hepatitis C. An acute infection is typically mild, but once chronic and untreated, may cause life threatening conditions, including liver cirrhosis and hepatocellular carcinoma. Approximately 70 million people worldwide live with chronic hepatitis C, with 400,000 related deaths annually [10]. Notably, hepatitis C can be cured in more than 95% of cases with direct acting antivirals that inhibit viral replication [10].

The re-emerging dengue virus (DENV) is a *Flavivirus* and belongs, as HCV, to the family *Flaviviridae.* Annually, DENV infects 390 million people worldwide, with around 96 million of them becoming symptomatic. Unlike HCV, DENV is vector-borne and is spread mainly by the mosquitoes of the *Aedes* species. Infection with DENV causes flu-like illness, occasionally with severe complications mostly associated with heterotypic secondary infections (e.g. hemorrhagic fever and shock syndrome) [11]. The clinical manifestation of a DENV infection is closely related to infections with the mosquito-borne chikungunya and Zika virus, leading to frequent misdiagnosis [12].

Coxsackieviruses are members of the genus *Enterovirus* of the family *Picornaviridae.* This genus includes important human pathogens such as poliovirus, enterovirus-A71 (EV-A71), EV-D68, coxsackievirus, and rhinovirus. Enteroviruses cause 10 to 15 million infections every year and therefore belong to the most prevalent pathogens [13]. Enteroviruses cause a variety of diseases, including hand-foot-and-mouth disease, encephalitis, meningitis, and paralysis [14]. Coxsackie B viruses are also known to infect cardiac tissue, leading to viral myocarditis, which can develop to congestive heart failure [15]. In this study, we focus on coxsackievirus B3 (CVB3).

Despite their broad range of clinical manifestations, transmission routes, and tropism (Table 1), plus-strand RNA viruses share remarkable similarities in their replication strategy. By definition, the genome of plus-strand RNA viruses has the polarity of cellular mRNAs. Therefore, after delivery into cells, the genome is directly translated, giving rise to a polyprotein that must subsequently be cleaved into viral proteins. These proteins induce host cell membrane rearrangements forming replication organelles (ROs). Either within those ROs or on its outer membrane facing the cytosol, viral RNAs are amplified by the viral replicase complex comprising, amongst others, the RNA-dependent RNA polymerase (RdRp). These ROs are thought to serve hiding viral RNAs from host immune response and thus to protect them from degradation. In addition, the membranous compartment allows the coordinated coupling of different steps of the viral replication cycle, i.e., RNA translation, RNA replication, and virion assembly [16–19].

However, there are striking differences in the viral life cycles of the three studied viruses. For example, the morphology of ROs in which replication takes place differs considerably. While HCV forms double membrane vesicles (DMV), DENV induces invaginations of host cellular membranes [20]. CVB3 infection first results in single-membrane tubular structures that subsequently transform into DMVs and multilamellar vesicles [21]. Additionally, HCV and DENV as representatives of *Flaviviridae* remodel membranes of the rough endoplasmic reticulum (rER), however, the *Picornaviridae* CVB3 uses the ER and Golgi apparatus for its RO formation [20]. Another interesting feature of CVB3 is its ability to trigger a so-called host translational shut-off, leading to increased viral over host RNA translation[22]. Repressed host RNA translation has also been reported for DENV [23], however, a host shut-off has not been reported for HCV, which instead shows parallel translation of viral and host cell RNAs, consistent with the predominantly chronic infection caused by this virus [24].

To identify an efficient, broadly active treatment strategy against viral infectious diseases, a comprehensive knowledge of viruses as well as their exploitive interaction with the host is of major importance. Mathematical modeling has proven to be a powerful tool to study viral pathogenesis, transmission, and disease progression and has increased our knowledge about therapeutic intervention and vaccination as well as the involvement of the immune system for viruses such as the human immunodeficiency virus (HIV), HCV, influenza A virus, DENV, Zika virus, and SARS-CoV-2 [25–31]. One of the major strengths of mathematical models is their ability to describe and analyze viral replication in a quantitative, dynamic (time-resolved) framework, and to characterize the influence individual parameters have on the ensuing dynamics. These models thus permit much deeper insights into viral replication and antiviral strategies than static, often more qualitative snapshots of host-pathogen interactions.

In the current study, we reproduced the dynamics of the initial post infection phase of the life cycle of three representative plus-strand RNA viruses, namely HCV, DENV, and CVB3, with one common mathematical model. Using the model, we identified pan-viral similarities and virus-specific differences in the life cycle of plus-strand RNA viruses that are represented by a unique set of model parameters. The inter-viral differences among the plus-strand RNA viruses under investigation have been further analyzed to study how these differences might be related to clinical disease manifestation, particularly with regard to chronic versus acute infections. Our model suggests that the number of ribosomes available for viral RNA translation may be a crucial factor for either acute or chronic infection outcome. Furthermore, we studied broad-spectrum antiviral treatment options and found inhibiting viral proteases involved in polyprotein cleavage, and RNA synthesis are promising drug targets.

## Methods

### Kinetic experiments and infectivity titers

#### HCV infections

2×10^5^ Lunet-CD81_high_ [49] cells per 6-well were seeded in 2 mL 16 hours prior to infection. To ensure simultaneous infection of all cells, cells were kept at 4°C for 30 min before medium aspiration and inoculation with pre-cooled PEG-precipitated HCV_cc_ (Jc1) [50] at an MOI of 1 at 4°C for one hour (1 mL per 6-well). The inoculum was removed and cells were covered with 1 ml per well pre-warmed (37°C) medium and incubated for one hour at 37°C. Medium was aspirated and cells were treated with an acid wash protocol to remove extracellular vesicles and unbound virus particles: cells were washed with an acidic solution (0.14 M NaCl, 50 mM Glycine/HCl, pH 3.0, 670 μL per 6-well) for three minutes at 37°C before neutralization with neutralization buffer (0.14 M NaCl, 0.5 M HEPES, pH 7.5, 320 μL per 6-well) and one wash with pre-warmed medium. After that, fresh medium was added. After indicated time-points, total cellular RNA was extracted by phenol-chloroform extraction. Infected cells were washed prior to lysis according to the acid wash protocol described above. After three washing steps with cold 1x PBS, cells were lysed in GITC buffer (700 μL per 6 well) and RNA was extracted as described [51]. A strand-specific RT-qPCR protocol was used to quantify numbers of (+)- and (-)-strand RNA per cell [52]. TCID50 of supernatants was measured and calculated as described previously [50] and converted to PFU/mL.

#### CVB3 infections

CVB3 wild-type (wt) and CVB3-Rluc, which carries *Renilla luciferase* upstream of the P1 region, were generated as described previously [53]. Subconfluent monolayers of HuH7 cells, provided by prof. R. Bartenschlager, were infected with CVB3 wt or CVB3-Rluc at an MOI of 1 for 45 minutes. After removal of the viral inoculum, cells were washed once with PBS and fresh medium (DMEM supplemented with 10% FBS and penicillin and streptomycin) was added. Every hour up to 9 hours post-infection, cells were collected and subjected to various assays. Each assay was performed on three biological replicates. Cells were either frozen together with the medium, after which progeny virus titers were determined by endpoint titration by the method of Reed and Muench and converted to PFU/mL.

Another set of cells were lysed in buffer to determine the luciferase activity as a measure of viral protein translation as described previously [53]. Lastly, cells frozen after aspiration of the medium were used for total RNA isolation and quantification of the amount of viral RNA copies per cell with quantitative PCR as described previously [54].

#### DENV infections

DENV kinetic measurements of intracellular plus-strand RNA and luciferase activity as well as extracellular infectious virus titers have been taken from [55]. In brief, 2×10^5^ Huh7 cells were infected with DENV reporter virus expressing Renilla luciferase [56] at an MOI of 10. RNA extraction and qRT-PCR as well as Renilla luciferase activity were analyzed from cell lysates. RNA was normalized to the 2 h value. Infectivity titers (TCID50/mL) were measured from viral supernatant by limited dilution assays and converted to PFU/mL, supernatants were subsequently supplemented [55].

### Plus-strand RNA virus replication model

We developed a mechanistic model using ordinary differential equations (ODEs) and mass action kinetics to analyze pan-viral similarities and virus-specific differences within the plus-strand RNA virus life cycle. Our published models on two plus-strand RNA viruses, HCV and DENV, served as a basis for the pan-viral plus-strand RNA virus replication model [19,55,57]. However, in our previous published models, we studied host dependency factors responsible for cell line permissiveness and restriction factors such as the innate immune response. Therefore, those models were modified to reflect merely the plus-strand RNA life cycle from virus entry to release of all viruses considered here.

The resulting model of plus-strand RNA virus replication is composed of four main processes: Entry of plus-strand RNA virus via receptor-mediated endocytosis and release of the viral genome (Fig 1 steps ① and ②), its subsequent translation into viral proteins (Fig 1 steps ③ to ⑤), viral RNA replication within the replication organelle (Fig 1 steps ⑥ to ⑨), and further replication (Fig 1 step ⑩) or RNA export out of the replication organelle (Fig 1 step ⑪) or virus packaging and release from the cell with subsequent re-infection of the same cell or infection of naïve cells (Fig 1 steps ⑫ and ⑬).

**Figure 1:**
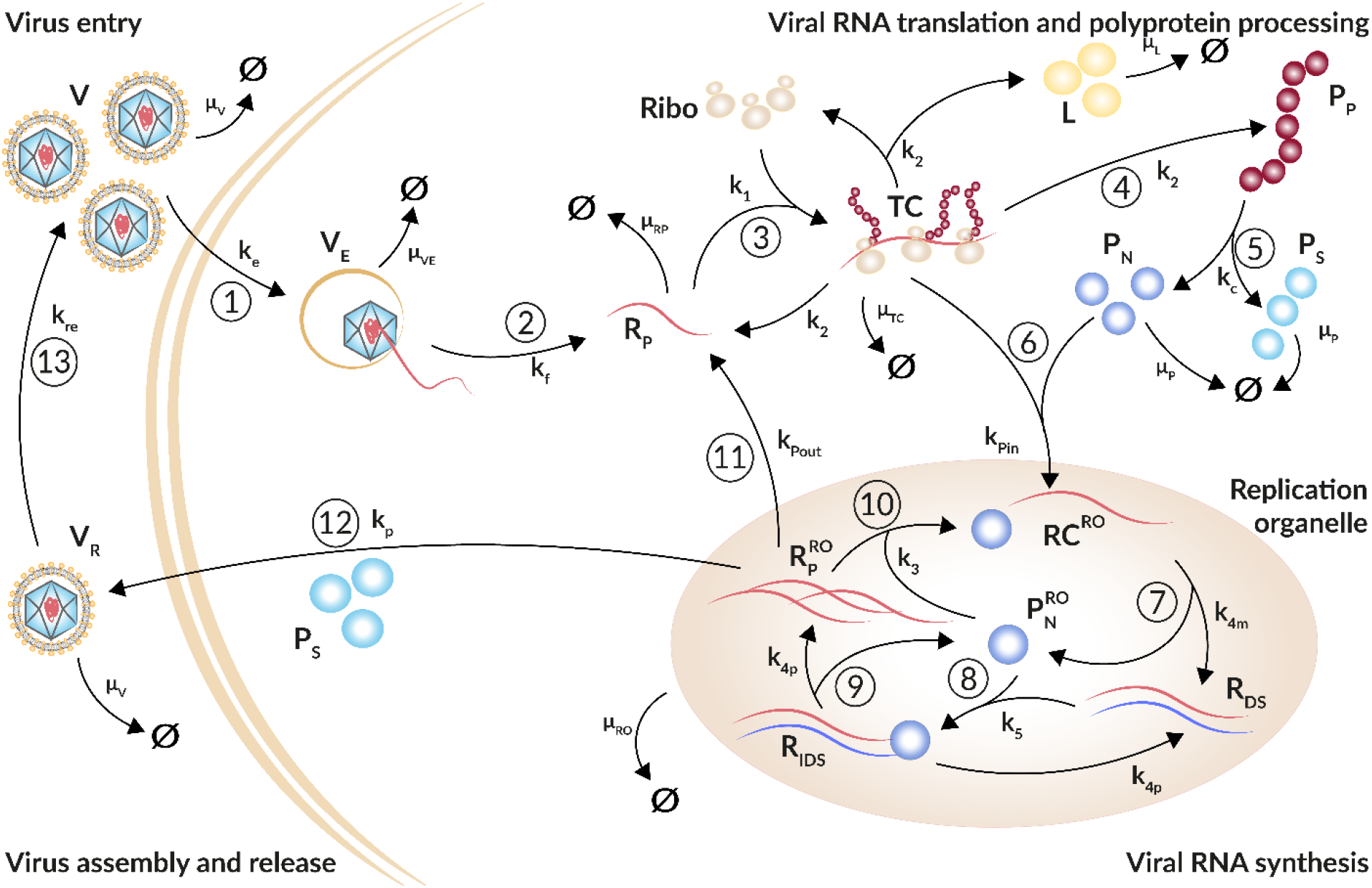
Schematic illustration of the plus-strand RNA life cycle. ① Virus (V) enters the cell via receptor-mediated endocytosis (k_e_). ② The viral genome (R_p_) is released (k_f_). Virus within the endosome (V_E_) degrades with rate constant *μ_VE_*. ③ Ribosomes (Ribo) bind the viral genome and form (*k*_1_) a translation initiation complex (TC) that degrades with rate constant μ_TC_. ④ The viral genome (R_p_) is translated (k_2_) into a polyprotein (P_p_) that ⑤ is subsequently cleaved (k_c_) into structural and non-structural viral proteins, P_s_ and P_N_, respectively. To measure translation activity, luciferase (L) is integrated into the viral genome and produced with RNA translation. Viral proteins degrade with rate constant μ_P_; luciferase degrades with rate constant μ_L_. ⑥ Non-structural proteins and freshly translated viral RNA form (k_Pin_) replicase complexes (RC) that are associated with replication organelles (ROs) and ⑦ serve as a template for the minus-strand synthesis (k_4m_) leading to double-stranded RNA (R_DS_).⑧ Viral non-structural proteins, such as the RdRp, within the replication organelle 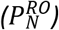 bind to doublestranded RNA forming (k_5_) a minus-strand replication intermediate complex (R_IDS_) that ⑨ initiates plus-strand RNA synthesis (k_4p_) giving rise to multiple copies of viral plus-strand RNA 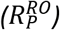. All species within the replication organelle degrade with the same rate constant μ_RO_. ⑩ The viral genome can remain within the replication organelle, where it undergoes multiple rounds of genome replication (k_3_), ⑪ it can be exported (k_Pout_) out of the replication organelle into the cytoplasm starting with the translation cycle again, or ⑫ the plus-strand RNA genome 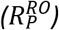 is packaged together with structural proteins (P_S_) into virions (V_R_) that are released from the cell (k_p_) and ⑬ may re-infect the same cell or infect naïve cells (k_re_). Extracellular infectious viral species (V and V_p_) degrade with rate constant μ_V_.

The virus infection process (Eqs. 1 and 2), i.e., receptor-mediated virus entry, fusion, and release of the viral genome into the cytoplasm, as well as re-infection of the same cell or further infection of naïve cells (Eq. 14) are represented by extracellular virus *V,* virus within endosomes *V_E_,* and newly produced virus released from infected cells *V_R_* and are given by the equations

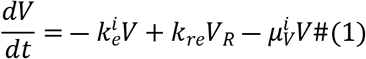

and

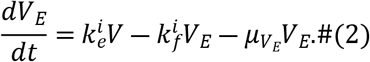

Extracellular virus *V* enters a single cell via receptor-mediated endocytosis with rate constant 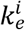 or degrades with constant rate 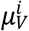. Note that virus-specific parameters are marked with a superscripted *í* with *i*∈{*HCV, DENV, CVB3*}. Virus within endosomes *V_E_* either degrades with rate constant *μy_E_* or undergoes conformational changes of its nucleocapsid resulting in the release of the viral genome *R_P_* with rate constant 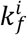. Note that extracellular virus is also replenished by the release of virus from the cell at rate *k_re_*.

Viral RNA translation and replication (Eqs. 3 to 13) are modeled based on our published HCV and DENV models [19,55]. In brief, our model describes the translation associated processes in the cytoplasm (Eqs. 3 to 8) starting with free viral RNA *R_P_* in the cytoplasm, an intermediate translation initiation complex *TC,* as well as the translated polyprotein *P_P_* which is cleaved into structural and non-structural viral proteins, *P_S_* and *P_N_*, respectively. Note that a firefly luciferase gene has been integrated into the viral genomes. The luciferase activity *L* was measured from cell lysates as a marker for translation activity (see Methods) reflecting protein concentration and has been introduced into the model. Translation and polyprotein processing are modeled with the following ODEs, where 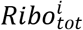 and *RC_MAX_* are the total number of ribosomes and maximal number of replicase complexes in a cell (see below for details), respectively:

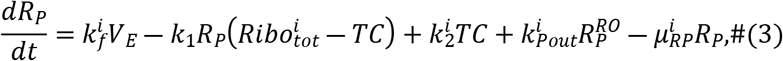

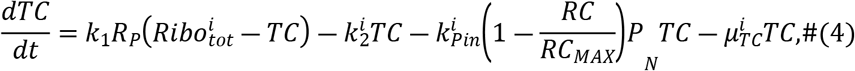

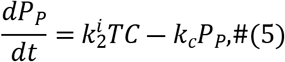

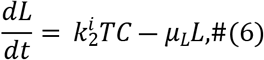

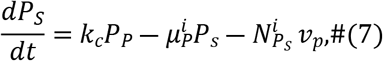

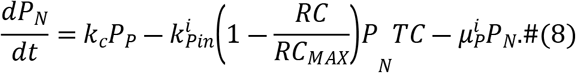

With rate constant *k*_1_ free host ribosomes form a translation complex *TC* with the viral plus-strand RNA genome *R_P_*. The total number of ribosomes 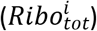 available for viral RNA translation was assumed to be constant and the number of free ribosomes is given by 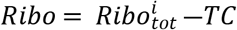. Note that 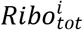 is only a fraction of the total cellular ribosome number. Translation of the viral plus-strand RNA genome generates the viral polyprotein *P_P_* and luciferase *L* with rate constant 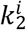. The viral polyprotein *P_P_* is subsequently cleaved with rate constant *k_c_* into structural and non-structural viral proteins, *P_S_* and *P_N_*, respectively. The translation complex *TC* decays with rate constant 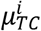, while luciferase and viral proteins degrade with rate constants *μ_L_* and 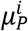, respectively. Note that for simplicity we assume structural and non-structural proteins degrade with the same rate constant, which has been summarized as one virus-specific viral protein degradation rate 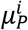.

The subsequent processes of viral RNA synthesis in the replication organelle (RO) are modeled by Eqs. 9 to 13 representing the replicase complex *RC,* double-stranded RNA *R_DS_*, a double-stranded RNA intermediate complex *R_IDS_*, newly synthesized viral plus-strand RNA in the RO 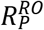, and non-structural proteins within the RO, 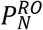, as follows:

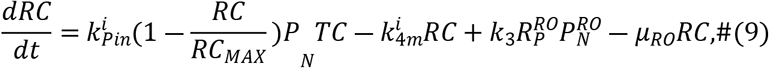

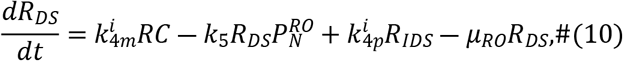

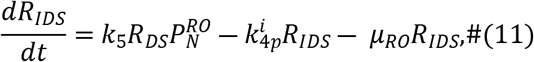

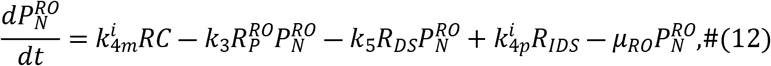

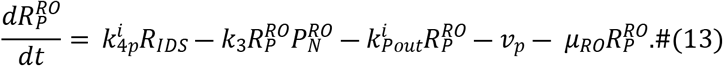

Viral non-structural proteins recruit the viral RNA after translation to the replicase complex [58]. Hence, for viral RNA synthesis, we require translated viral RNA, i.e., the translation complex *TC* instead of free cytosolic viral RNA *R_P_* to interact with the non-structural proteins. Thus, the translation complex *TC* together with a subset of non-structural proteins *P_N_* are imported into the RO, where they lead to the formation of a replicase complex *RC* with rate constant 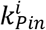. Following successful replicase complex formation, ribosomes dissociate from the complex as is accounted for in Eq. (4). We furthermore assume that there is a limitation in the number of replicase complexes formed within a cell. To do so, we extend 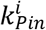 by 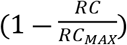 with the carrying capacity for replicase complexes *RC_MAX_* [57,59].

Within the RO, minus-strand RNA synthesis occurs from the replicase complex with rate constant 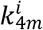, leading to the formation of double-stranded RNA *R_DS_*, which along with the non-structural proteins are released from the RO, 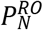. Subsequently, the double-stranded RNA binds again to 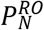 with rate constant *k*_5_ to form a double-stranded intermediate replicase complex *R_IDS_*, initiating plus-strand RNA synthesis with rate constant 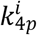. For simplicity, we assume that minus and plus-strand RNA synthesis occur with the same rate constant 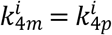. The newly synthesized plus-strand RNA genomes 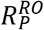 either remain within the RO to make additional replicase complexes with rate constant *k*_3_, are exported out of the RO into the cytoplasm for further RNA translation with export rate 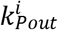, or are packaged together with structural proteins into virions *V_R_* and are subsequently released from the cell. Assembly and release of virus particles is represented by a Michaelis-Menten type function *V_p_* described below (Eq. 15, compare [55,60]). The RNA and protein species within the RO (*RC, R_DS_, R_IDS_*, 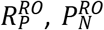) are assumed to degrade with the same decay rate *μ_RO_* and represent the decay of the entire replication organelle.

The released virus *V_R_* may re-infect the same cell or infect new cells with rate constant *k_re_*, or degrade with rate constant 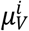, resulting in the equation

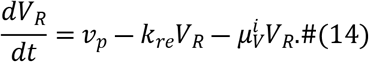

Assembly of newly synthesized viral plus-strand RNA genome 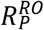 and viral structural proteins *P_S_* into viral particles and their subsequent release from the host cell are described using a Michaelis-Menten type function, with rate

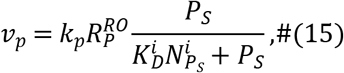

where *k_p_* is the virion assembly and release rate and 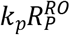 being the maximum release rate that is limited by viral resources. Let 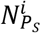 be the number of structural proteins in a virus of type *i*, then to produce virus at rate *v_p_* will require a large number of proteins 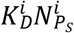, where 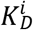 is a scaling constant and 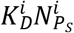 is the number that corresponds to the half-maximal release rate [see [55,60,61] for more details].

### Pan-viral and virus-specific model parameters

To complete the model of the plus-strand RNA virus life cycle, we need to specify model parameters. To prevent overfitting and parameter uncertainty, we fixed many parameter values to either experimentally determined values or to values estimated in other modeling studies. In some cases, we were able to calculate rate constants directly, such as for viral RNA translation and synthesis, which could thus be fixed as described in S1 Supporting text. An overview of all parameters values is given in Table 2.

**Table 2:** Parameter values and 95% confidence intervals in (). Note that parameter values marked with * were fixed due to previous assumptions and calculations. Furthermore, confidence intervals marked with + hit the set estimation boundary; ± calculated from the data; # experimentally measured for Zika virus; ≠ experimentally measured for poliovirus.

| Parameter | Description | HCV | DENV | CVB3 | Unit |
| --- | --- | --- | --- | --- | --- |
| $k_e^i$ | Virus entry rate | 10 (1.9, $10^+$) | 0.31 (0.28, 0.34) | 1.3 (0.9, 1.7) | 1/h |
| $k_f^i$ | RNA release rate | 10 (1.7, $10^+$) | 0.008 (0.006, 0.01) | 0.016 (0.006, 0.04) | 1/h |
| $k_1$ | Formation rate of the translation complex | 1000 (840, $1000^+$) | | | mL/molecule h |
| $k_2^i$ | Virus RNA translation rate | 180 [65] | 100 [55] | 300 <sup>‡</sup> [66] | 1/h |
| $k_c$ | Polyprotein cleavage rate | 2.24 (1.18, 7.4) | | | 1/h |
| $k_3$ | Formation of additional replicase complexes within the replication organelle | 42 (5.5, 525) | | | mL/molecule h |
| $k_{4m}^i = k_{4p}^i$ | Minus- and plus-strand RNA synthesis rate | 1.1 [65] | 1.0 [55] | 50 <sup>‡</sup> [66] | 1/h |
| $k_{pin}^i$ | Formation rate of the replicase complex | 4.4 (2.4, 7.5) | 0.45 (0.29, 0.74) | 1.4 (0.52, 4.09) | mL/molecule h |
| $k_5$ | Formation rate of the replication intermediate complex | 6018 (1549, 68401) | | | mL/molecule h |
| $k_{pout}^i$ | Export rate of viral RNA out of the replication organelle | 33 (0.8, 1477) | 53 (16, 432) | 0.23 (0.16, 0.43) | 1/h |
| $k_p$ | Assembly and release rate | 158 (47, $1000^+$) | | | mL/molecule h |
| $k_{re}$ | Reinfection rate | 0.01 ($0.01^+$, 0.038) | | | 1/h |
| $\mu_{RP}^i$ | Degradation rate of cytosolic viral RNA | 0.26 [65] | 0.23 [67] | 0.15 <sup>‡</sup> [68] | 1/h |
| $\mu_{TC}^i$ | Degradation rate of the translation complex | 0.13 * | 0.115 * | 0.075 * | 1/h |
| $\mu_{RO}$ | Degradation rate of viral RNA and protein within the replication organelle | 0.086 [19] | | | 1/h |
| $\mu_P^i$ | Degradation rate of viral protein | 0.08 [19] | 0.46 [67] | 0.43 [69] | 1/h |
| $\mu_L$ | Degradation rate of luciferase | 0.35 [19] | | | 1/h |
| $\mu_V^i$ | Degradation rate of extracellular infectious virus | 0.1 [57] | 0.13 [70] | 0.08 [71,72] | 1/h |
| $\mu_{VE}$ | Degradation rate of intracellular virus within the endosome | | 0.23 # [73] | | 1/h |
| $V_0^i$ | Initial virus concentration | 0.2 (0.16, 0.25) | 1 (0.8, 1.3) | 1 (0.4, 2.2) | molecules/mL |
| $Ribo_{tot}^i$ | Total ribosome concentration | 0.005 (0.004, 0.007) | 0.48 (0.41, 0.55) | 6.7 (5.0, 9.1) | molecules |
| $RC_{MAX}$ | Maximum number of replicase complexes | | 0.46 (0.34, 0.64) | | molecules/mL |
| $K_D^i$ | Scaling constant for virus | 0.04 ± | 1.8 ± | 40 ± | virions |
| $N_{P_S}^i$ | Number of structural proteins needed to produce 1 virion | 180 [65,74] | 180 [55,74] | 60 [15] | molecules/virion |
| $f_{R_P}^i$ | Scale factor for plus-strand RNA | 394 (274, 524) | 0.76 (0.58, 1.0) | 550 (245,1366) | |
| $f_{R_M}^i$ | Scale factor for minus-strand RNA | 1377 (945, 1872) | - | - | |
| $f_L^i$ | Scale factor for luciferase | - | 0.41 (0.33, 0.5) | 0.08 (0.06, 0.1) | |

### Parameter estimation, model selection, and model analysis

Our model has 61 parameters; 30 of them were fixed, while 31 were estimated by fitting the model to experimental data. As the fixed parameter values were experimentally measured, calculated, or taken from literature, we had information about which were virus specific (S1 Supporting text and Table 2). To determine which of the remaining model parameters are conserved across the different viruses considered (pan-viral) and which parameters are virus-specific, we performed several rounds of model evaluation using the Akaike information criterion (AIC) and model identifiability analysis (profile likelihood estimation). See S2 Supporting text for a description of the model selection process.

We fit the plus-strand RNA virus replication model simultaneously to the virus-specific data sets for HCV, DENV, and CVB3. To fit the mathematical model to the experimental data, we calculated the total plus-strand RNA 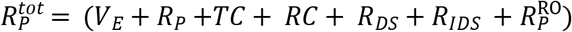, total minus-strand RNA 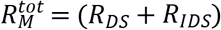, luciferase *L*, and total infectious virus *V^tot^* = (7 + *V_R_*). Note that our model accounts for infectious virus since infectious titers were measured for all three viruses. Further note that for the infectious virus measurements for HCV, *V^tot^* = *V_R_*, since measuring infectious virus started 20 h pi. We introduced three scale factors *f_L_*, *f_R_M__*’, and *f_R_P__* to re-scale experimental measurements acquired in relative measurements (plus-strand RNA for DENV), molecules per cell (plus-and minus-strand RNA measurements for HCV and plus-strand RNA for CVB3) and relative light units (luciferase for DENV and CVB3).

We implemented the model in MATLAB (The MathWorks) 2016 using the Data2Dynamics toolbox [62]. We assessed model identifiability using the profile likelihood estimation method implemented in Data2Dynamics [62,63]. In Data2Dynamics, a parameter is identifiable if its 95% confidence interval is finite [62,63]. Note that an estimated model parameter may hit a predefined upper or lower parameter boundary which hampers the calculation of the 95% confidence interval. In such cases, a one-sided 95% confidence interval has been calculated starting from the estimated model parameter and thus with its upper or lower boundary marked with + in Table 2. Details about the model fitting and model selection process are in S1 Supporting material.

We performed a global sensitivity analysis in MATLAB using the extended Fourier Amplitude Sensitivity Test (eFAST) [64]. We calculated sensitivities with regard to the total plus-strand RNA 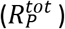 concentrations throughout the course of infection. We studied hypothetical drug interventions by including the effects of direct acting antivirals (DAA) into the model. For this purpose, we simulated putative drugs targeting (1) viral entry and internalization *k_e_*, (2) release of the viral RNA genome *k_f_*, (3) formation of the translation initiation complex *k*_1_, (4) viral RNA translation *k*_2_, (5) polyprotein cleavage *k_c_*, (6) replicase complex formation *k_Pin_*, (7) minus- and plus-RNA synthesis *k*_4*m*_ and *k*_4*p*_, as well as (8) virus particle production and release (*v_p_*). To introduce drug effects into the model, we assumed a drug efficacy parameter 0 ≤ *ε* ≤ 1, and multiplied the parameters above by (1 – *ε*) to simulate drug treatment. Similar to our previously published DENV model, we calculated the average virus particle concentration released from the cell upon drug administration (*ε* ≠ 0) until 5 days post drug administration, i.e., a drug treatment observation window of 120 h. The average virus particle concentration with treatment (*ε* ≠ 0) has been normalized to the average virus concentration without drug treatment (*ε* = 0). Note that we studied two different time points of drug administration: at the very beginning of the infection, 0 h pi, and when the system is in steady state, 100 h pi.

## Results

As shown in Fig 2 (left panels), the model replicates the experimental data for all three viruses. The comparison of their plus-strand RNA and virus (infectious particles) dynamics, reveals virus-specific characteristics. CVB3 is fast-replicating with a life cycle of about 8 hours (depending on the cell type) after which the infected cells begin to die. Similarly, DENV is also cytopathic but seems to be slower replicating and thus has a longer life cycle than CVB3 with infectious particles being produced at about 16 h pi [56]. In contrast, HCV is non-cytopathic with a much longer life cycle. In our experimental measurements, the CVB3 viral load peaked at 8 h pi with 193 PFU/mL/cell. The HCV viral load peaked with 0.06 PFU/mL/cell around 44 h pi, while the DENV viral load reached its maximum with approximately 8 PFU/mL/cell around 10 hours earlier at 30 to 34 h pi (Fig 2A, 2B, 2C). We calculated the corresponding average virus concentration per measurement time point for HCV, DENV, and CVB3 per cell as 0.04 PFU/mL/cell, 1.8 PFU/mL/cell, and 40 PFU/mL/cell, respectively. Thus, the average infectious HCV viral load was only 4% of the average DENV viral load and only 0.3% of the average CVB3 viral load. Similarly, CVB3 reached a peak of almost 500,000 plus-strand RNA copies per cell at 8 h pi, while HCV produced only 10,000 copies per cell at 70 h pi, i.e., 98% less than CVB3.

**Figure 2:**
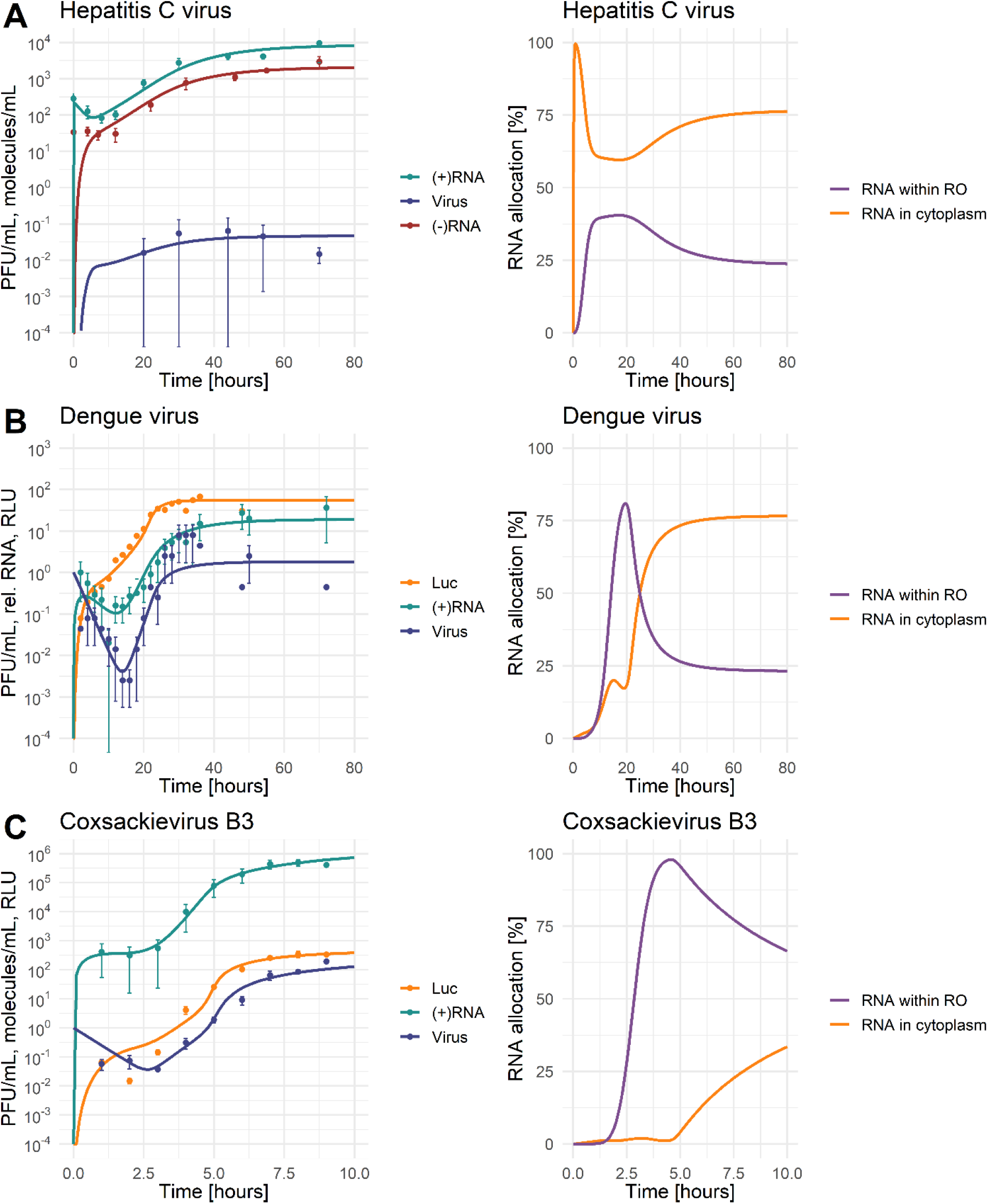
Best model fit (solid line) to the data with standard deviation (left) and model prediction of plus-strand RNA allocation between cytoplasm and replication organelle (RO) (right). For parameter values see Table 2. [LEFT: 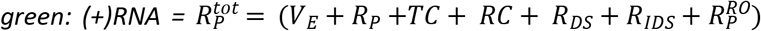, red: 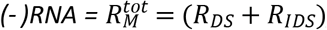, blue: A) and B) Virus = V^tot^ = (7 + 7R) or C) Virus = V^tot^ = V_R_, yellow: Luc = L; RIGHT: yellow: RNA in cytoplasm 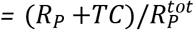, purple: RNA within replication organelle 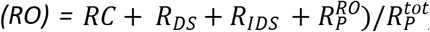; Infectious virus in PFU/mL, (+) and (-)RNA were measured in molecules/mL or relative RNA concentration, luciferase was measured in relative light unit (RLU)]

### Model selection and uncertainty

The intracellular model structure has been taken from our previously published HCV model [19], upon which we built with our recently published DENV model [55]. However, a striking difference from our previous HCV and DENV models is the absence of host factors involved in replicase complex formation and/or virus assembly and release. We have previously shown that host factors are recruited by the virus and seem to be beneficial for host cell permissiveness and virus replication efficiency [19,55]. Instead, here we describe inter-viral replication differences with virus-specific parameter sets based on model evaluation by AIC and profile likelihood estimation (see Methods, S1 and S2 supporting texts).

Including the maximal number of replicase complexes (*RC_MAX_*) improved the basic model AIC from 3025 to 1982 and thus served as a starting point for the virus specific model selection process (see S1 Supporting material). After several rounds of model selection by comparing AICs and taking model identifiability into account, we added five virus specific processes to our basic model (from a total of 13 considered processes): (1) the total number of ribosomes 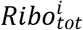 available for viral RNA translation, (2) virus entry 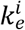, (3) viral genome release 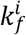, (4) formation of the replicase complex 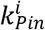, and (5) export of viral RNA from the RO into the cytoplasm 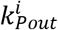. Note that based on literature data and previous assumptions, we fixed some virus-specific and pan-viral processes and degradation rates (see S1 Supporting text and Table 2). The best-fit model showed high similarity to the virus-specific experimental measurements and a high degree of model identifiability (see Fig 2 for best fit, Fig 3 for the parameter profiles based on the profile likelihood estimation, and Table 2 for parameter values with 95% confidence intervals).

**Figure 3:**
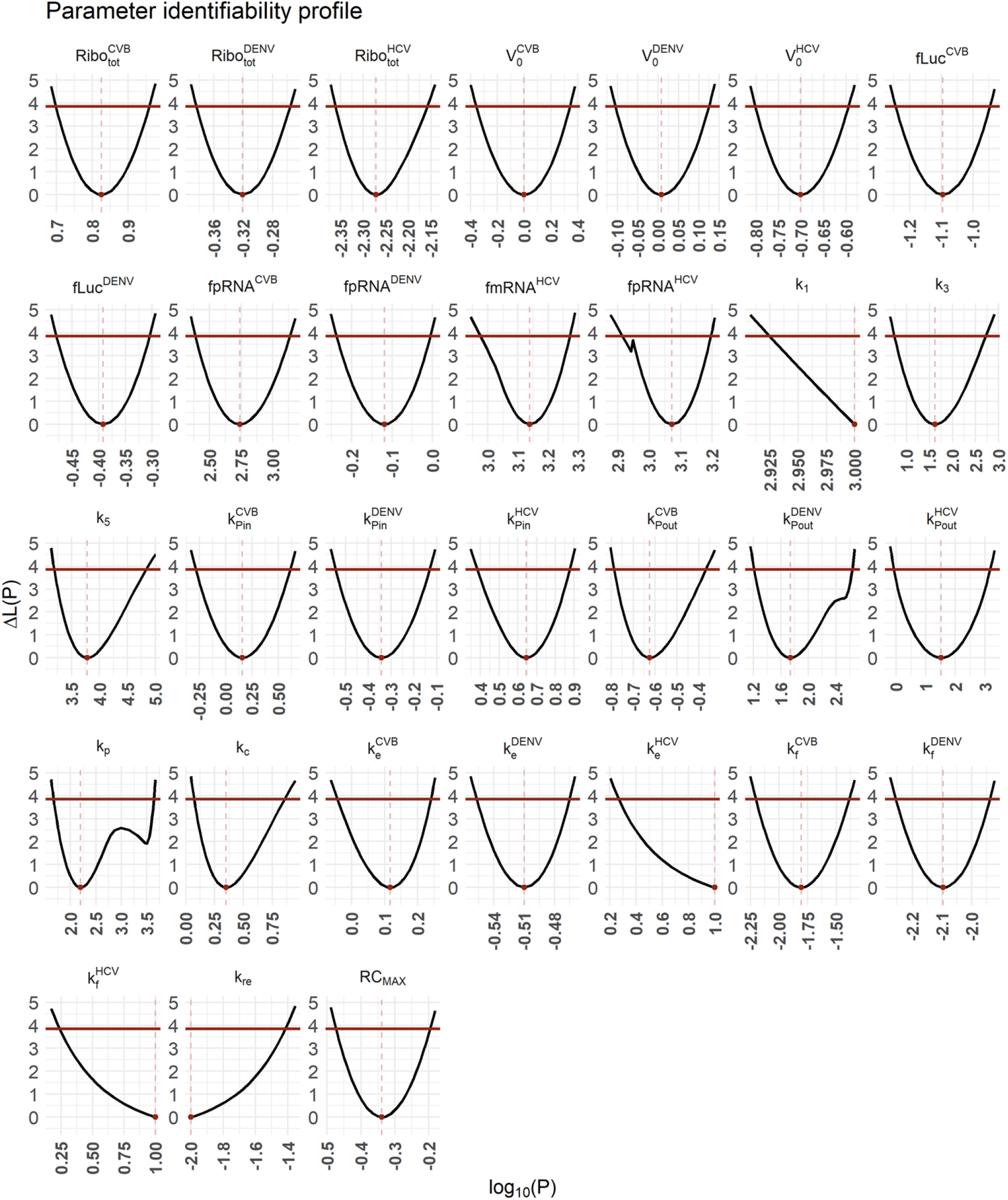
Uncertainty analysis of the best-fit model. For parameter values and 95% confidence intervals see Table 2. The best fit is shown in Fig. 2.

### RNA allocation

The allocation of plus-strand RNA in the cytoplasm and within the RO, as predicted by our model, shows interesting virus-specific differences (Fig 2 right panel). Compared to the total amount of viral RNA, HCV has most of the RNA allocated to the cytoplasm and thus available for viral RNA translation at any given time. In DENV, our model predicted that the allocation strategy changes throughout the viral life cycle, with the majority of plus-strand RNA within the RO initially. At around 25 h pi, viral RNAs are equally distributed between the two compartments, while at the end of the DENV life cycle the majority of viral RNA is in the cytoplasm. Interestingly, at steady state, the predicted allocation of both HCV and DENV is the same, with 25% of RNA allocated to the RO and 75% to the cytoplasm. In contrast, the predicted viral RNA allocation is opposite for CVB3. CVB3 has the majority of RNA available within the RO, which contributes to the 2 to 3 log higher viral load.

### Virus specificity

For a successful virus infection, the first hurdles to overcome are virus entry and the release of the viral genome into the cytoplasm. The rate constants for virus entry 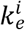 and vRNA release 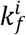 had the highest estimated values for HCV. However, both values were practically non-identifiable suggesting a limitation in the amount of data. Hence, we could only estimate the lower boundary of the 95% confidence intervals, which suggest 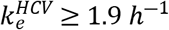 and 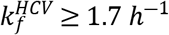. CVB3 seems to be slightly better adapted to the cell line with a 4-times higher entry rate and 2-times higher vRNA release rates compared to DENV. According to our model selection process, the degradation rate of internalized virus within endosomes *μvE* was pan-viral suggesting neither an advantage nor disadvantage for the studied viruses.

The next processes in the viral life cycle are vRNA translation and polyprotein processing with parameters *k*_1_ for the formation of the translation initiation complex, 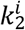 vRNA translation, and *k_c_* polyprotein cleavage. Models including virus-specific *k*_1_ or *k_c_* either did not improve the quality of the model fit (no AIC improvement) or were non-identifiable when tested as virus-specific and thus have been selected as pan-viral (see S2 Supporting material). However, the viral RNA translation rate 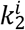 was calculated based on genome size and ribosome density and set as virus-specific (see S1 Supporting text). In the vRNA translation and polyprotein processing step, the only parameter our model selected as virus specific was the total number of ribosomes 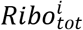. Since the ribosome number has been selected in the first round of model selection (see S2 Supporting text), it emphasizes the importance of this host factor with CVB3 showing the highest estimated ribosome number available for RNA translation. In contrast, HCV and DENV use only 0.07% and 7% of the ribosomes CVB3 uses, respectively. Interestingly, increasing the number of ribosomes in the HCV life cycle to those of CVB3 (from 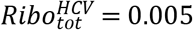 to 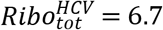 molecules per ml) increases the infectious virus load by three orders of magnitude (Fig 4A). In the same way, decreasing the number of ribosomes in the CVB3 life cycle to those of HCV (from 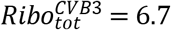 to 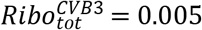 molecules per ml) decreases the CVB3 virus load by three orders of magnitude (Fig 4B). In contrast, when increasing the viral RNA synthesis rates of HCV to those of CVB3 (from 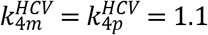 to 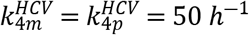), the viral load did not increase. However, decreasing the viral RNA synthesis rates of CVB3 to those of HCV (from 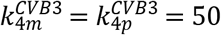 to 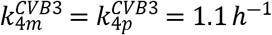) decreased the viral load by one order of magnitude. This suggests an important role of ribosomes as key players in the production of structural and non-structural proteins necessary for efficient vRNA replication and virus production.

**Figure 4:**
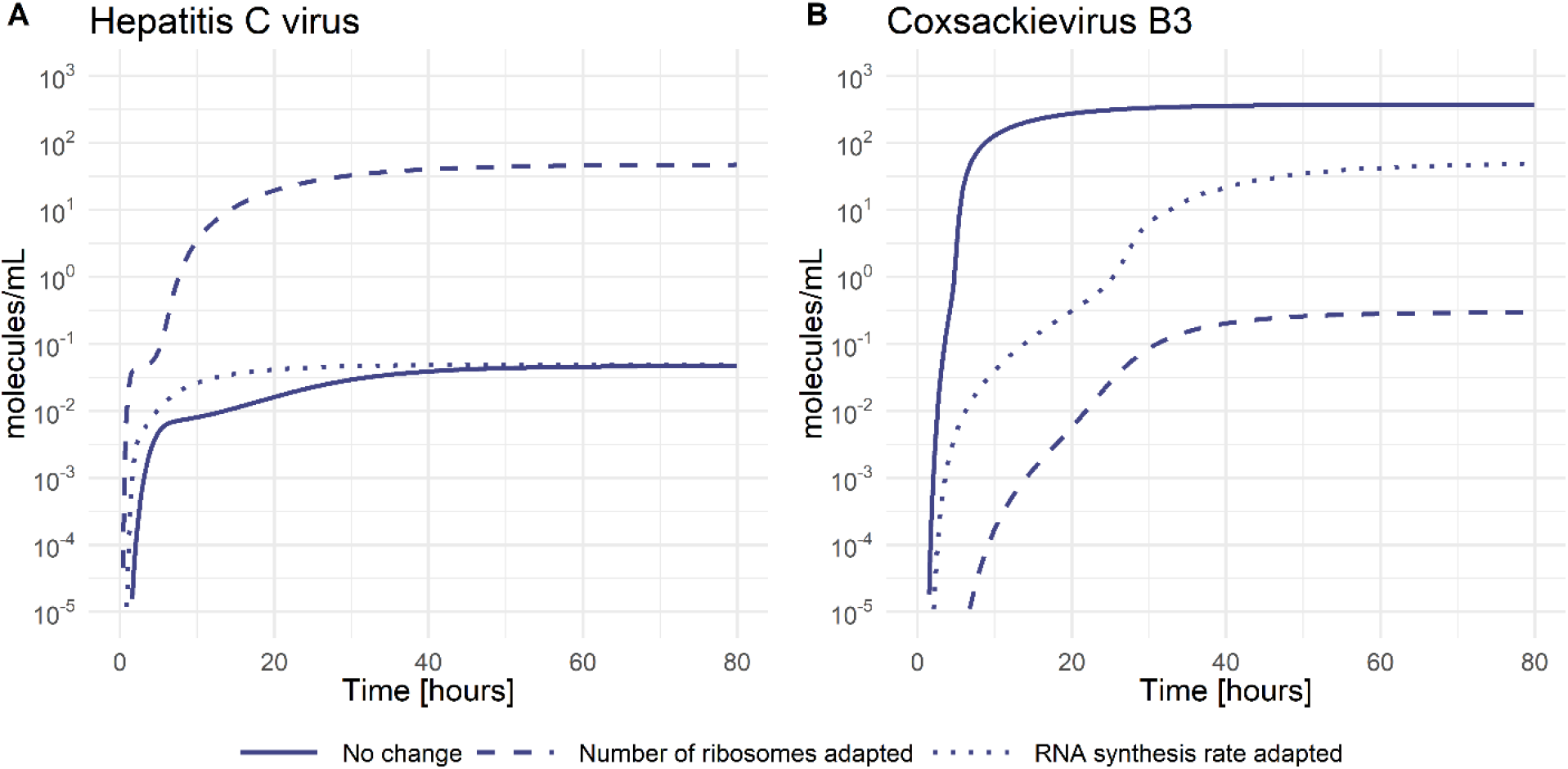
Infectious virus concentration with parameter adjustments. **A)** HCV concentration with estimated parameters (solid), the number of ribosomes taken from CVB3 (dashed), and the RNA synthesis rate taken from CVB3 (dotted). **B)** CVB3 concentration with estimated parameters (solid), the number of ribosomes taken from HCV (dashed), and the RNA synthesis rate taken from HCV (dotted).

The subsequent processes of vRNA replication depend on successful viral protein production. Viral non-structural proteins are crucial for the formation of the replicase complex and its formation rate 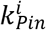, which has been selected as virus specific. Here, HCV seems to be more efficient and better adapted to the Huh7 cell line, showing a 10- and 4-times faster formation rate compared to DENV and CVB3, respectively. Furthermore, our estimated replicase complex formation rates suggest that the formation of double membrane vesicles may be more efficient (HCV and CVB3) compared to the formation of invaginations (DENV). However, the maximum number of replicase complexes *RC_MAX_* as well as the degradation of species within the RO (*μ_RO_*) were not selected as virus-specific, especially since the viral RNA synthesis rates were initially set as virus-specific (Table 2). Interestingly, even though being a pan-viral model parameter, not all viruses reached the maximal number of replicase complexes *RC_MAX_* (the carrying capacity). The dynamics of replicase complexes shows a clear separation between DENV and CVB3 versus HCV (Fig. 5A and 5B). CVB3 reached the estimated carrying capacity around 5 h pi, while DENV reached 98% of the possible carrying capacity around 25 h pi. Strikingly, the replicase complex formation for HCV reached its maximum at a 74% lower level of the pan viral carrying capacity, even though our model estimated the fastest RC formation rate for HCV.

**Figure 5:**
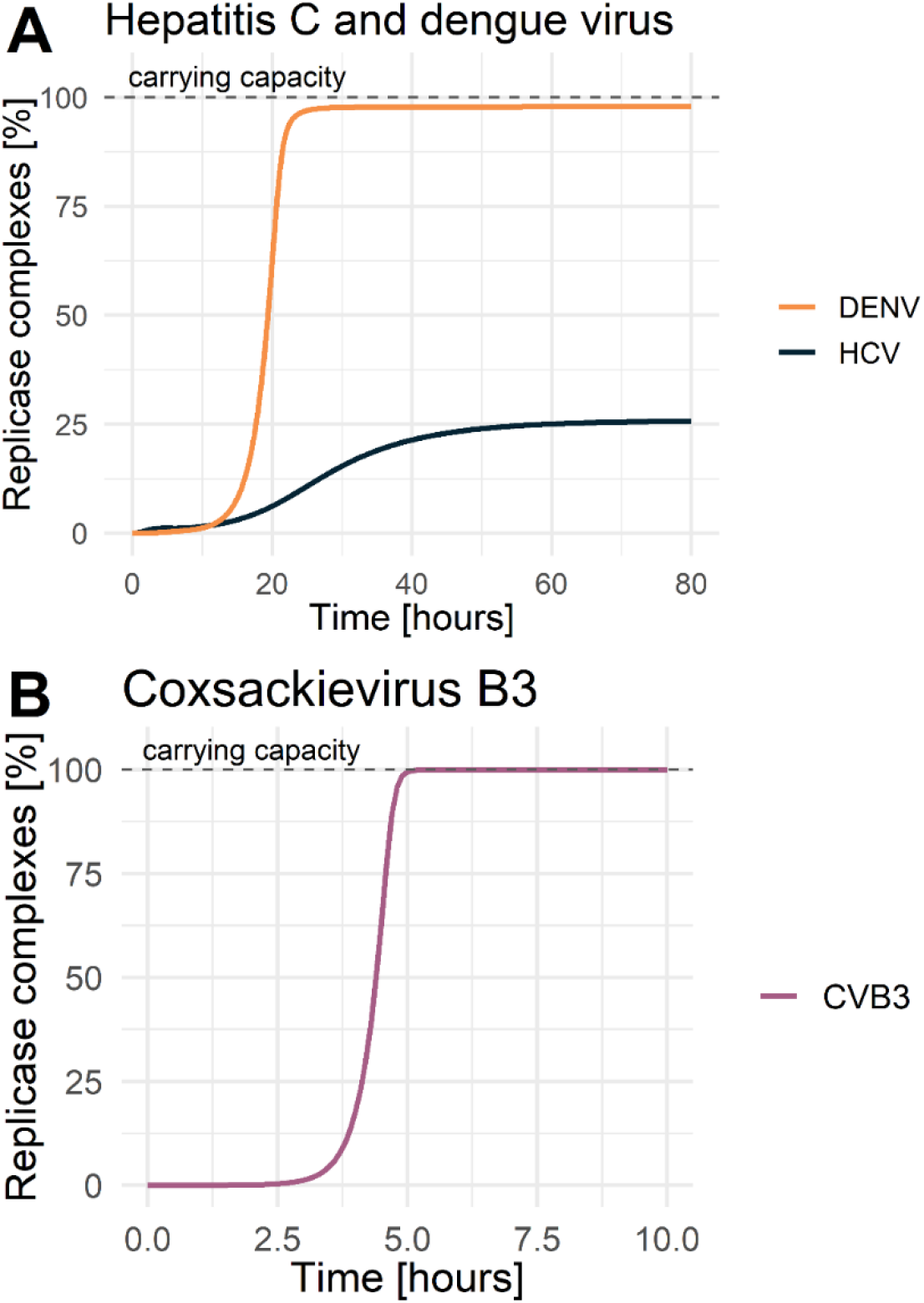
Replicase complexes over time. Dynamics of replicase complexes for **A)** hepatitis C and dengue virus, **B)** coxsackievirus B3. The dashed grey line represents the carrying capacity or the maximum number of formed replicase complexes.

The export of viral RNA from the RO to the site of RNA translation 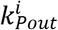 has also been selected as virus specific, where HCV and DENV seem to be more efficient than CVB3 which showed an almost 190 times slower trafficking process.

Following the production of viral proteins and RNA genomes, the single components assemble into virions and are released from the cell. Here, the virus assembly and release rate *k_p_* as well as the reinfection rate *k_re_* have been selected as pan-viral, while the scaling constant 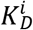 as well as the number of structural proteins necessary per virion 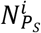 were calculated from the data or taken from the literature, respectively, and thus set as virus-specific (Table 2).

### Sensitivity analysis and drug intervention

Having a detailed model of the intracellular replication of plus-strand RNA viruses, we next addressed the question of which processes shared across all viruses showed the highest sensitivity index to potential drug interventions (Fig 6). Our sensitivity analysis suggests that model parameters associated with vRNA translation 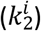 and synthesis within the RO (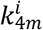 and 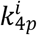) are highly sensitive for all viruses. Furthermore, all viruses were sensitive to the formation of replicase complexes 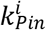 and its maximum number *RC_MAX_*. Interestingly, over the course of infection, DENV and CVB3 showed a time-dependent sensitivity pattern beginning with viral entry 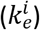 being sensitive, followed by the release of the viral genome 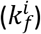. However, both model parameters were not sensitive for HCV, possibly due to practical nonidentifiability (see above). Moreover, vRNA translation and replication seem to start around 5 or 20 h pi in CVB3 and DENV, respectively, suggesting viral entry as a rate limiting process.

**Figure 6:**
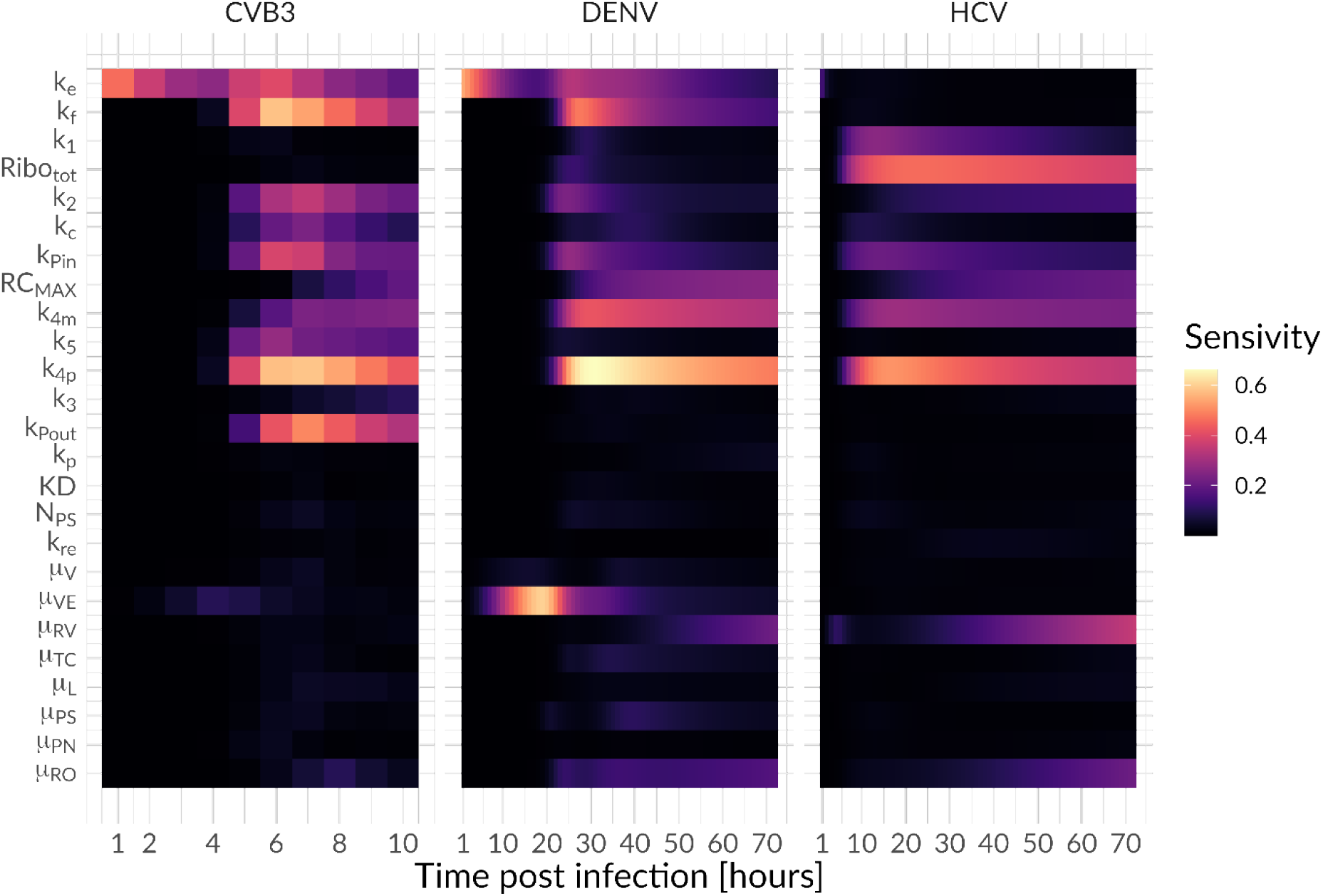
Global sensitivity profile for the model species plus-strand RNA over the course of infection (CVB3 = 10 hours, HCV = DENV = 72 hours).

There are also some interesting differences between the three viruses. While the formation of the translation initiation complex (*k*_1_) showed a higher sensitivity in HCV, vRNA translation 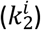 was more sensitive for CVB3 and DENV. Furthermore, for HCV, the number of ribosomes available for HCV RNA translation was one of the most sensitive parameters, while having negligible sensitivity for CVB3 and DENV. This may be a reflection of the strength of the IRES (CVB3) or the 5’ UTR/Cap (for DENV), where a strong IRES may require less ribosomes for robust recruitment to initiate vRNA translation. However, for CVB3 viral RNA export 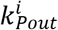 is among the most sensitive processes, while being not sensitive for HCV and DENV. Interestingly, the degradation of virus in endosomes (*μ_VE_*) showed the highest sensitivity among the degradation rates for DENV early in infection (around 10 to 25 h pi), while the degradation of cytosolic vRNA (*μ_RP_*) seem to be highly sensitive towards the end of infection for both DENV and CVB3.

As a next step, we aimed to analyze if any processes can be targeted leading to a 99% reduction in extracellular virus upon inhibition. We therefore studied the effects of inhibiting core processes of the viral life cycle (Fig 7). We then simulated *in silico* the administration of a hypothetical drug at two different time points using our mathematical model: at the very beginning of the infection (0 h pi) or at steady state (100 h pi). For all viruses and both drug administration time points, we determined the critical drug efficacy, *ε,* where the viral life cycle is successfully inhibited and the *in-silico* infection is cleared. Note that we define a virus infection as being cleared if extracellular virus is reduced by more than 99%. By testing both drug administration time points, we found that at the beginning of infection (0 h pi) inhibiting any process led to an eradication of the virus (Fig 7). Since the viral replication machinery is not established, viral entry and vRNA release may be possible drug targets, however, an almost 100% inhibition (*ε*~1) was necessary to block the infection process (S1 Table). Obviously, *in-silico* drugs targeting virus entry and vRNA release at a time point after an established viral infection, is not able to reduce the viral load. However, for both drug administration time points, targeting vRNA translation as well as vRNA synthesis showed the strongest effect, and thus are the most promising drug targets (S1 Table). Interestingly, targeting the formation of the replicase complexes could not clear (or even reduce) CVB3 infection with a drug administration given at steady state (S1 Table). Moreover, in the case of DENV, targeting vRNA export from the RO into the cytoplasm at steady state led to a 6% increase in virus with incomplete inhibition. Only a 100% inhibition and thus a drug efficacy of 1 was able to clear the virus by 99%.

**Figure 7:**
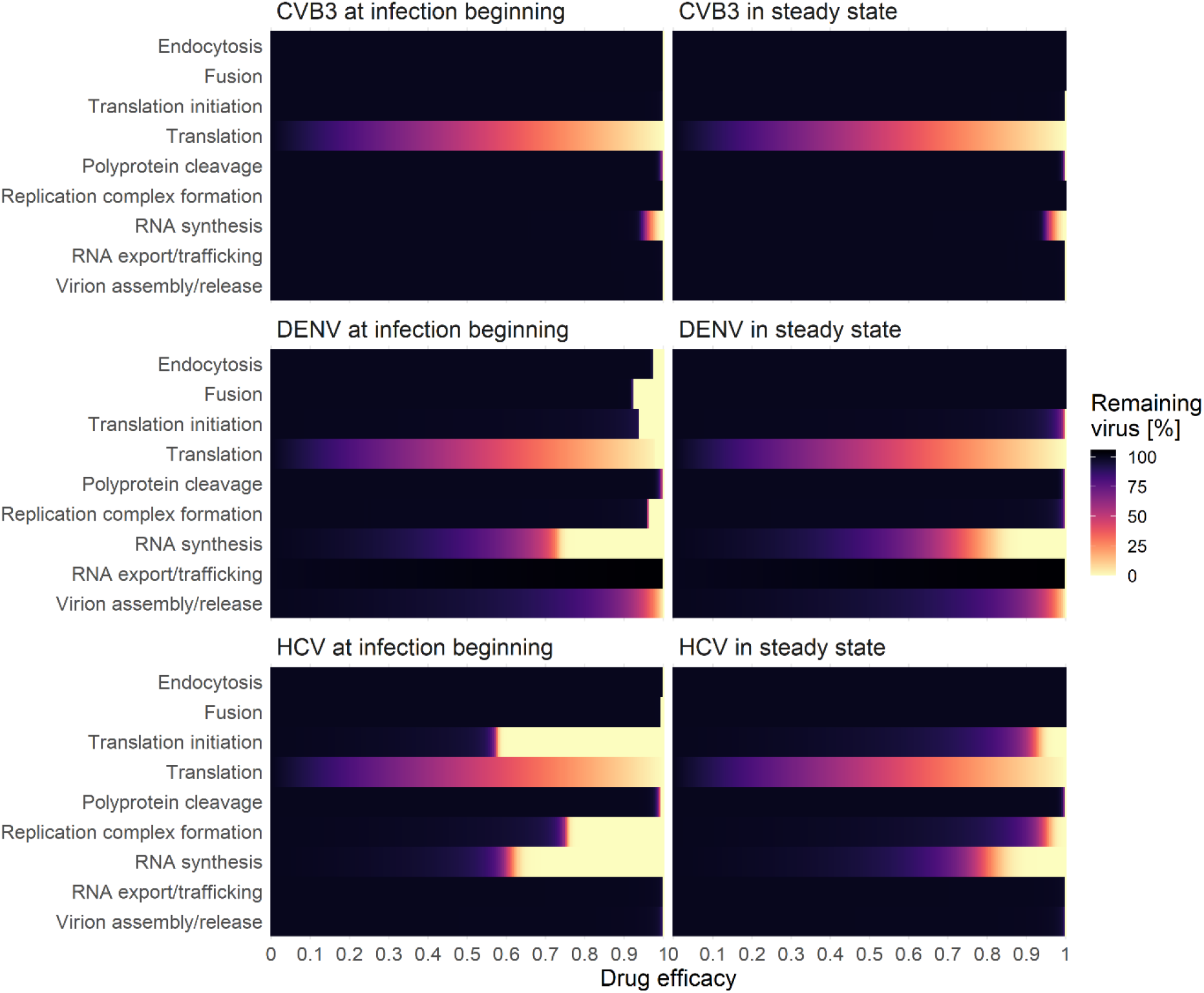
Effects of drug interventions at two different time points: at infection beginning (left) and in steady state (right). A successful drug treatment leads to a more than 99% viral eradication (light yellow), while an ineffective drug treatment leads to 100% remaining virus (black).

Since most direct acting antiviral drugs are highly efficient in combination, we determined the critical drug efficacy of individual drugs inhibiting either translation complex formation, vRNA translation, or polyprotein cleavage used in combination with drugs that inhibit vRNA synthesis or formation of the replicase complex at steady state (Figs 8 and 9 and S1 and S2, Figs, S1 Table). We identified the “sweet spot” for efficient viral eradication (by more than 99%). Our model predicted that HCV and DENV showed a comparable pattern of viral clearance to a combination of two drugs, while for the clearance of CVB3 higher drug efficacies were necessary to clear the infection. Inhibiting vRNA synthesis in combination with vRNA translation or polyprotein cleavage by more than 90% was an efficient combination for HCV and DENV (Fig 8B and 8C, S1 Table, S2A Fig). However, to clear the infection in all viruses, vRNA synthesis and translation or polyprotein cleavage, have to be inhibited by more than 99% or 98%, respectively (Figs 9B and 9C). Interestingly, inhibiting vRNA synthesis and translation complex formation by more than 76% showed the overall lowest critical drug efficacy to clear an HCV infection. Nevertheless, for CVB3, the vRNA synthesis and translation complex inhibition need to be higher than 99.3% to clear the infection with an almost 10 hours delay in viral clearance (Figs 8A and 9A, S1 Table). Overall, we found the lowest pan-viral critical drug efficacy was for the combined inhibition of vRNA synthesis and polyprotein cleavage with a required 98% effectiveness for each drug (Figs 8C and 9C, S1 Table,). Note that we also tested *in silico* the combination therapy of inhibiting translation complex formation, vRNA translation, and polyprotein cleavage together with replicase complex formation. However, higher critical drug efficacy constants were needed to clear the infection (S1, S2 Figs and S1 Table).

**Figure 8:**
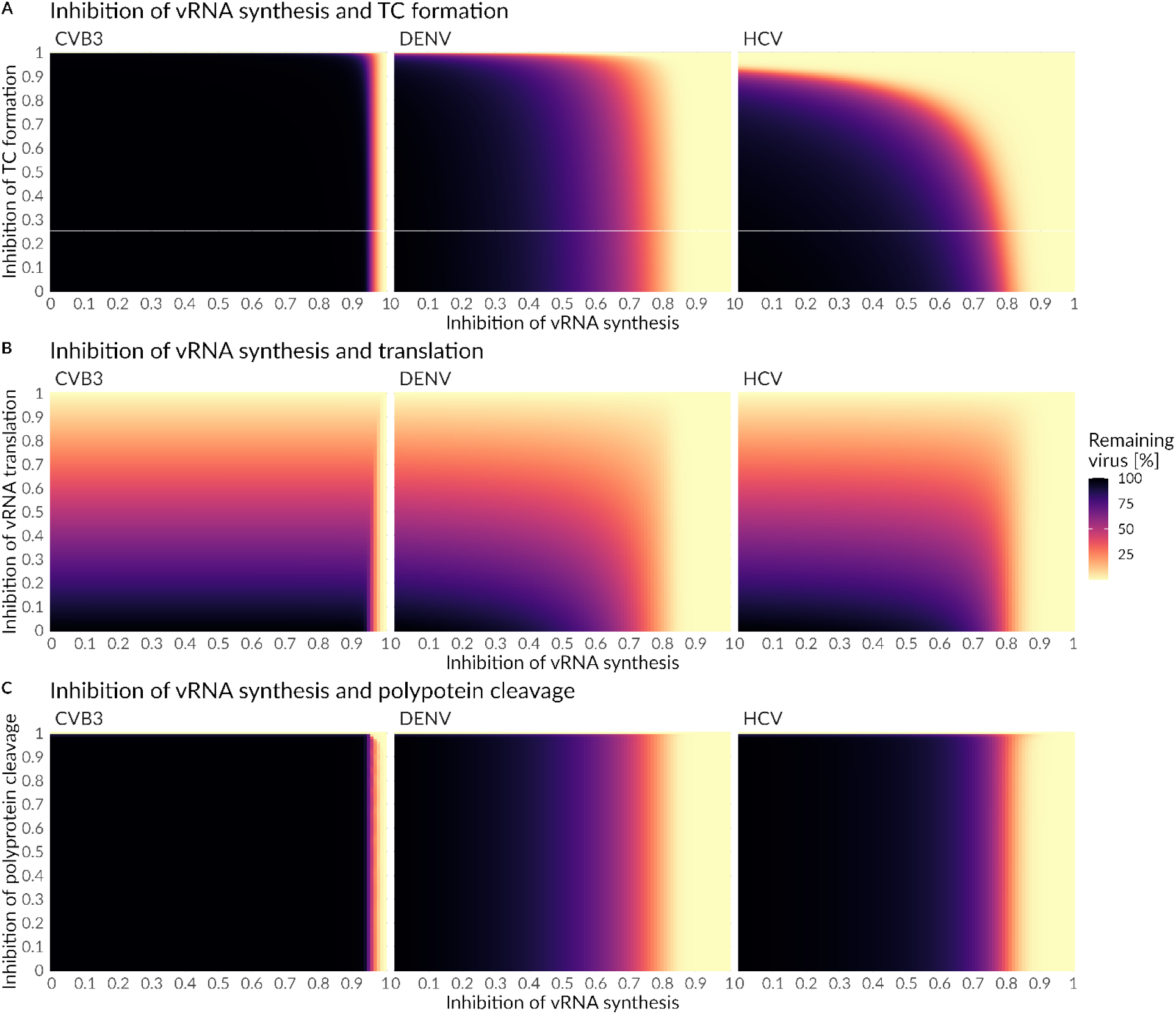
Combined drug effects on **A)** vRNA synthesis and formation of translation complex (TC), **B)** vRNA synthesis and translation, and **C)** viral RNA synthesis and polyprotein cleavage. Initiation of treatment was in steady state (100 h pi). A successful drug treatment leads to more than 99% viral eradication (light yellow), while an ineffective drug treatment leads to 100% remaining virus (black).

**Figure 9:**
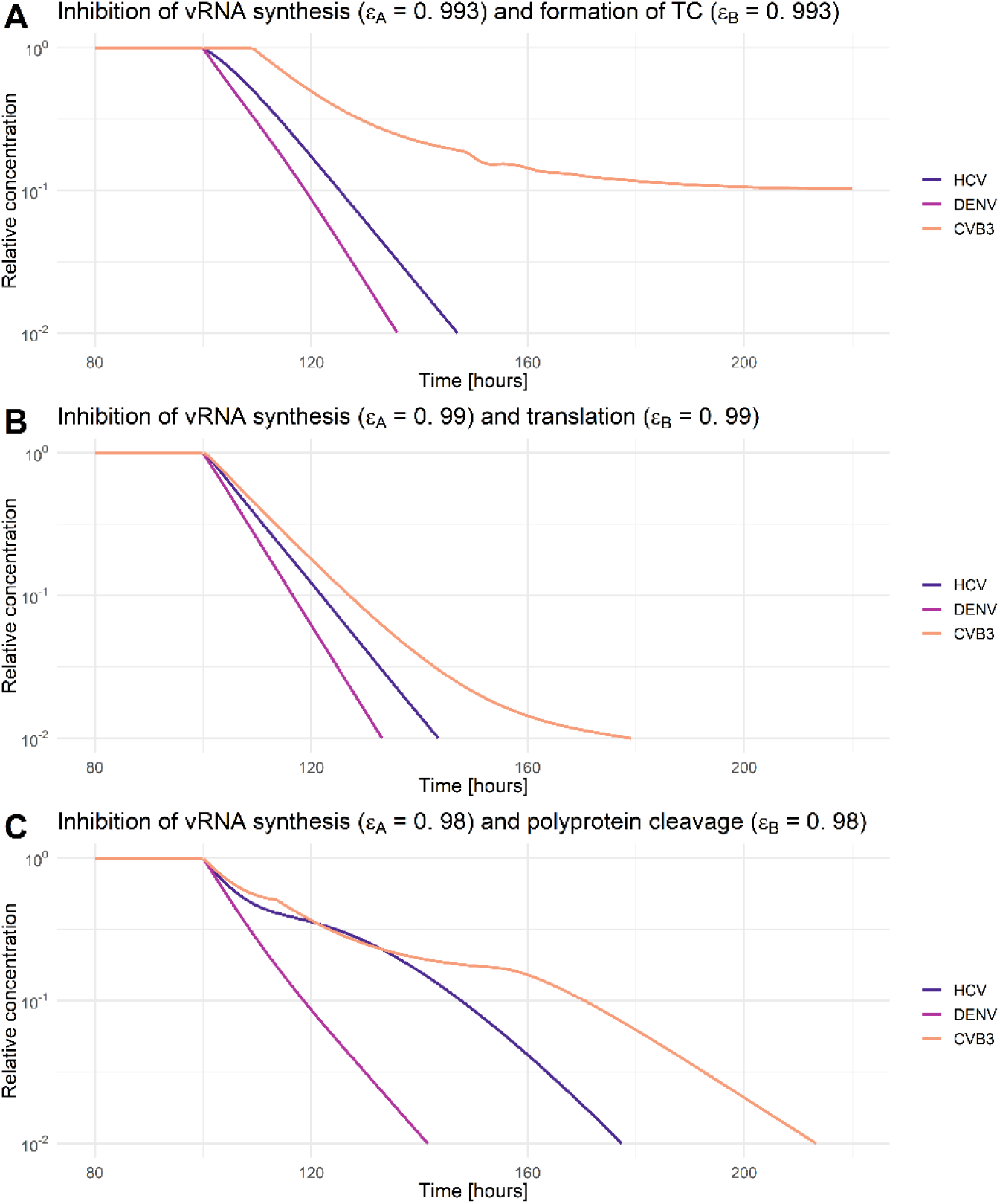
Relative virus decay under combination therapy that clears HCV, DENV, and CVB3 infections. A combined drug effect on **A**) vRNA synthesis and formation of translation complex (TC), **B)** vRNA synthesis and translation, and **C)** viral RNA synthesis and polyprotein cleavage. Initiation of treatment was in steady state (100 h pi). The drug efficacy constant (ε_A_ and ε_B_) were chosen as minimal efficacies to clear all three viruses. For comparability, virus-specific concentrations in steady state have been normalized to their virus-specific pre-treatment steady state concentration. A successful drug treatment leads to a more than 99% viral eradication (light yellow), while an ineffective drug treatment leads to 100% remaining virus (black).

## Discussion

Mathematical modeling of viral dynamics has a long history and has been applied to a variety of viral infectious diseases [25]. Population based models considering susceptible and infected cell populations, especially studying virus-host interactions and treatment opportunities for HIV, HCV and Influenza, represent the most prominent mathematical models in the field [25,75–78]. However, mathematical models considering intracellular viral replication mechanisms in detail are still limited and are usually developed for one specific virus such as HCV [19,57,59,79,80], DENV [55], HIV [81], or influenza A virus [60,61,82–87]. Recently, Chhajer et al (2021) studied with a simplified mathematical model the viral life cycles of the plus-strand RNA viruses HCV, Japanese encephalitis virus, and poliovirus. The authors mainly focused on the slow and delayed kinetics of the intracellular formation of replication organelles, which may predict infection outcome [88]. To our best knowledge, we present here the first mathematical model that studies simultaneously the complexity of intracellular viral replication kinetics for three different representatives of plus-strand RNA viruses, namely HCV, DENV, and CVB3, measured in the same cell line – Huh7. The basis for our present study were our previously published intracellular models for HCV [19,57] and DENV [55], which we generalized and adapted to reflect the intracellular replication mechanisms of plus-strand RNA viruses more broadly, as well as the underlying experimental conditions. We compare viral replication mechanisms as well as pan-viral similarities and virus-specific differences, which may help to understand acute or chronic infection outcome that in turn may be an initial step towards the development of broad-spectrum antiviral treatment strategies.

Our best-fitting model showed high similarity with the virus-specific data and a high degree of parameter identifiability. However, it showed one shortcoming in capturing the dynamics of the experimental measurements of virus in DENV: the viral peak and subsequent drop of the extracellular DENV concentration around 32 h pi. However, in our previously published DENV model, we showed that the dynamics of extracellular infectious virus was dependent on host factors that were packaged into the virions [55]. Since we did not include host factors into the current model, except for ribosomes, our aim was to describe the average extracellular virus dynamics for the first 25 h pi. In the final model, we estimated 31 parameters of which 27 were identifiable. The 95% confidence intervals of four parameter values hit the upper or lower boundary of estimation, where changing of the parameter boundaries by up to 1000-fold did not lead to an improvement of the model fit or to improved identifiability.

The non-identifiable rate constant of the naïve cell infection *k_re_* may be explained by the fact that reinfection in our culture system may not occur for each virus. However, the process remained in the final model because of different MOI infection experiments, where a lower MOI (MOI of 1 as in the case of CVB3 and HCV) may account for multiple rounds of infection. The formation rate of the translation initiation complex *k*_1_ seems to be a non-identifiable process in the model structure, as it was also non-identifiable in our previous DENV model [55]. Further, the model processes of virus entry and vRNA genome release, *k_e_* and *k_f_*, were practically non-identifiable for HCV. A possible explanation for both processes being non-identifiable may be insufficient experimental measurements for HCV to uniquely estimate both rate constants, e.g., the lack of intracellular protein concentration measurements for HCV. However, since both parameters were identifiable for CVB3 and DENV and both processes were selected as virus-specific, 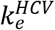 and 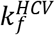, they remained virus-specific in the final model.

### Virus specific differences and pan-viral similarities

Studying similarities and differences in the viral RNA translation and replication strategies of different viruses is experimentally challenging. Our mathematical model may help to shed light on this topic by studying 25 processes from cell infection to release of the newly packaged infectious virions. Five processes within the viral life cycle were determined to be virus-specific: (i) virus entry, (ii) release of vRNA genome, (iii) the number of ribosomes available for vRNA translation, (iv) formation of replicase complexes, and (v) trafficking of newly produced viral genomes from the RO into the cytoplasm.

#### Virus internalization and genome release

The three viruses we studied each have different internalization processes mediated by differences in attachment/entry versus uncoating receptors [89]. HCV replicates *in vivo* in hepatocytes and consequently HCV showed the most efficient internalization and genome release processes in our studied hepatocyte derived Huh7 cells. *In vitro,* HCV replicates most efficiently in Huh7 cells and its closely related sub-clones, while the infection of other cell lines has been challenging [90]. However, both DENV and CVB3 have a broad tropism. DENV infects monocytes, macrophages, and dendritic cells and CVB3 infects brain and cardiac tissue as well as hepatocytes [15,35,91–93]. Thus, the faster internalization and genome release of CVB3 in comparison to DENV, and thus its ability to replicate very well in Huh7 cells, is not surprising due to its broader cellular tropism.

#### Viral RNA translation

Among the plus-strand RNA viruses we studied, CVB3 represents the fastest replicating virus with a life cycle of around 8 to 10 hours. Newly synthesized CVB3 RNA is detectable at two h pi in the Golgi apparatus, the site of ROs and thus vRNA synthesis. Levels of viral RNA increase rapidly and peak four h pi [94]. One key feature of successful CVB3 RNA replication is its ability to shut off host mRNA translation, carried out by the virus by degrading eukaryotic initiation factor eIF4G important for the cellular cap-dependent translation complex formation. The result is not only the rapid availability of non-structural proteins required for replicase complex formation [95], but also a lower level of components of the cell’s intrinsic immune response. Interestingly, we found the highest total ribosome availability for CVB3, in agreement with its ability to shut-off the translation of the host’s mRNA while keeping vRNA translation high due to a very efficient internal ribosome entry site (IRES). According to our calculated viral RNA translation rate constants, translation is 2 to 3 times faster compared to HCV and DENV, respectively. It has been shown that the polysome size – the number of ribosomes bound to a single CVB3 RNA molecule, which translate the viral genome at the same time – is around 30 ribosomes per polysome, but changes over the course of the CVB3 life cycle; 40 ribosomes per polysome at the beginning of the CVB3 life cycle and 20 ribosomes later in infection [66,96]. Furthermore, Boersma et al. (2020) found that CVB3 translation rates were independent of host translation shup down. However, the authors speculated that a host translation shut down may boost the CVB3 translation at the end of its life cycle where host cell resources may be limited [97]. Conversely, for DENV it has been shown that the DENV RNA template is only sparsely loaded with ribosomes and showed a low translation efficiency [98]. Nevertheless, Roth et al. (2017) found that the host’s mRNA translation decreases during DENV infection, suggesting that DENV also has the ability to repress the host mRNA translation although not as efficiently as CVB3 [23]. A partial host cell RNA translation shut-off and consequently a higher number of ribosomes available for DENV RNA translation is predicted by our model, with DENV having the second highest predicted ribosome concentration. Interestingly, even though DENV is able to partially shut down the host’s mRNA translation, this suppression does not seem as efficient compared to the complete CVB3 host shut-off.

#### Formation of the replicase complex

Our model suggests a faster formation of double membrane vesicles compared to invaginations, i.e., HCV and CVB3 showed faster replicase complex formation compared to DENV. Compared to DENV and CVB3, HCV showed a 10- and 4-times faster rate of replicase complex formation, respectively. A possible reason may be cell tropism with hepatocellular-derived Huh7 cells being the cell line of choice for studying HCV. Interestingly, the host mRNA translation shut-off of CVB3 was not associated with a faster supply of non-structural proteins (RdRp) and thus faster replicase complex formation. However, host cell translation shut off may be associated with higher availability and more efficient utilization of viral resources for the formation of replicase complexes, as suggested by our model. CVB3 reached the maximal number of replicase complexes after around 5 h pi, while HCV used 76% less of the possible cell’s carrying capacity. However, cell tropism and thus a specific set of host factors involved in the process of replication organelle and replicase complex formation may be the crucial factors in this process, as we have shown previously for HCV and DENV [19,55].

#### Viral RNA export from the RO into the cytoplasm

A striking difference between *Flaviviridae* (HCV and DENV) and *Picornaviridae* (CVB3) concerns the parameter values and model sensitivity against changes of the trafficking of newly synthesized vRNA from the RO to the site of translation. For CVB3, our model suggests intra-compartment trafficking two orders of magnitude slower as compared to HCV and DENV, with a highly significant sensitivity of this parameter against changes. A possible explanation may lie in the involvement of different compartments or cell organelles in vRNA translation and replication. All viruses need close proximity to the rough endoplasmic reticulum and its ribosomes for successful vRNA translation; however, they use different cytoplasmic membranes and thus different sites for the formation of their ROs and thus for vRNA synthesis. *Flaviviridae* remodel mainly the rough endoplasmic reticulum, using membrane vesicles or invagination as the site for vRNA translation and synthesis without being exposed to the (possibly damaging) cytoplasmic environment. Melia et al (2019) found that CVB3 uses the rough endoplasmic reticulum first and the Golgi later in infection, suggesting a high degree of flexibility and adaptation of CVB3 to its environment. To what extent viral replication occurs on either membrane is unknown, however, other studies suggest that Golgi-derived membranes serve as the main origin of viral replication [94,99,100]. During CVB3 infection, the Golgi collapsed and was not detectable anymore, suggesting that ROs were Golgi derived [101]. Regarding efficient viral protein production for virion packaging, CVB3 is not enveloped and may only need a fraction of the structural proteins that DENV and HCV needs for assembly (see S1 Supporting text for details), implying that CVB3 developed strategies to overcome longer trafficking distances. However, another explanation may be a possible regulation and competition of vRNA translation and virion packaging. Early in infection, vRNA may be used for translation, while later in infection vRNA may be packaged into virions and thus not available for vRNA translation.

### Hypothetical mechanisms behind acute and chronic infections

The plus-strand RNA viruses studied here share the major steps in their life cycle and their replication strategy, but despite these similarities show very different clinical manifestations. While HCV has a relatively mild symptomatic phase, it can establish a chronic infection with low-level viral replication over decades, that goes mostly undetected by the host’s immune response. In contrast, DENV causes a vigorous acute self-limited infection that can become life-threatening. Similarly, CVB3 usually causes an acute infection with flu-like symptoms but can become chronic. The underlying mechanisms for the development of chronic infections are unclear, our plus-strand RNA virus replication model might help to reveal the differences in the viral dynamics leading to different clinical manifestations.

DENV/ZIKV and CVB3 produce a higher ratio of plus- to minus-strand RNA (20:1) compared to HCV, with a plus- to minus-strand RNA ratio of 3:1 (measured in our data) up to 10:1 (reported in literature [102–109]), which may be HCV-strain or cell line-specific. One may speculate that a higher viral RNA synthesis rate may be responsible for the higher plus-to minus-strand RNA ratio in viruses causing acute infections. However, our calculated vRNA synthesis rates were comparable for HCV and DENV, but 50 times lower compared to the CVB3 RNA synthesis rate which may be due to faster vRNA copying or faster *de novo* initiation of vRNA synthesis. In HCV, studies found an RNA synthesis rate of 150 to 180 nt/min [110,111], however, the rate of RNA synthesis in DENV is to our knowledge unknown. Nevertheless, Tan et al. (1996) found low in vitro polymerase activity for DENV NS5, which is in line with the polymerase activities for West Nile and Kunjin viruses, suggesting that this is a conserved feature of flavivirus polymerases [112] and possibly *Flaviviridae* including HCV.

As for CVB3, it has been shown that the closely related PV synthesizes a single RNA template in 45 to 100 sec [66]. Additionally, it is estimated that between 3 and 10 RdRps are bound to one single PV RNA genome. However, in our plus-strand RNA model, we did not consider the RdRp density bound to one single viral RNA template, due to a lack of data for HCV and DENV. According to our model predictions, key processes for a faster viral life cycle may be a combination of: (1) faster viral RNA translation and synthesis rates and/or faster vRNA synthesis initiation, (2) host cell translation shut-off and thus higher ribosome availability for viral RNA translation and at the same time lower ribosome availability for antiviral protein production, (3) and shorter RNA half-lives for intracellular viral RNA (more important in cell lines with intrinsic immune responses or *in vivo).* Interestingly, the potential role of these key processes is in line with the results of the global sensitivity analysis: All CVB3 replication process rates within the RO show highly significant sensitivities, suggesting that CVB3 strongly depends on an efficient replicative cycle within the RO. Additionally, global sensitivities of vRNA degradation rates in the cytoplasm or within the RO seem rather negligible.

Our model predicted that an optimal usage of viral resources to form replicase complexes within a cell was only realized by DENV and CVB3. Strikingly, HCV only reached 26% of the cell’s replicase complex carrying capacity. A possible reason may be a limitation in viral resources to form replicase complexes such as viral RNA or non-structural proteins. Both may be again related to the lower availability of ribosomes for viral protein production in HCV, whereas DENV and CVB3 have the advantage of a partial or complete host cell translation shut off, respectively. However, virus-specific differences in the ribosome availability and translation activity may be related to different translation mechanisms. While HCV and CVB3 have IRESes, i.e., the RNA translation is cap-independent, DENV’s translation mechanism is cap-dependent. Furthermore, different IRES types have variations in their structural elements and recruit host factors as regulatory elements, which affects the translation initiation complex and viral RNA translation. Therefore, a higher ribosome availability for vRNA translation may be associated with different translation mechanisms such as different secondary structures and host factors assisting in ribosome binding [113–116]. Furthermore, a higher number of ribosomes available for vRNA translation may be directly associated with a higher production of viral proteins. However, the more ribosomes available for cellular mRNA translation and thus the production of proteins of the immune response, the higher may be the intracellular degradation of viral components, resulting in a limitation in viral resources. Ribosome availability and its control may thus be a crucial factor for viral replication efficiency.

To analyze this aspect further, we asked whether we could make virus production in HCV more efficient or CVB3 less efficient. Increasing the *in-silico* ribosome availability in HCV to that of CVB3 increased the viral load by three orders of magnitude. In contrast, a 50-fold increase in the HCV RNA synthesis rate had no effect on the viral load in steady state due to a limited availability of the viral RNA polymerase in the replication organelle [19]. In contrast, using only 0.07% of ribosomes for CVB3 RNA translation, thus setting the ribosome level to the number of ribosomes used in HCV, decreased the CVB3 viral load by three orders of magnitude. Interestingly, the coronaviruses nonstructural proteins, including those of SARS-CoV-2, target multiple processes in the cellular mRNA translation, causing a host cell translation shut off similar to CVB3 and DENV [117,118]. Therefore, a repression or complete shut-off of the host mRNA translation machinery may be a key-feature of acute viral infections.

Comparing *in vivo* viral dynamics with those of *in vitro* experiments is challenging. Nevertheless, we found comparable pattern of viral dynamics: reported *in vivo* and our *in vitro* experiments. *In vivo,* HCV showed an exponential growth rate of 2.2 per day [119], while DENV and CVB3 grow twice as fast with a rate of 4.3 and 4.5 per day in human and murine blood, respectively (approximated from [38,44]). However, in murine cardiac tissue, the *in vivo* CVB3 exponential growth rate increases to approximately 14.5 per day [38]. Furthermore, the different exponential growth rates are associated with variations in the peak viral load. At its peak, HCV produces 10^8^ RNA copies per g liver tissue [43], DENV produces 1 to 2 orders of magnitude more virus (10^9^ to 10^10^ RNA copies per ml blood) [44], and CVB3 produces 3 to 4 orders of magnitude more virus (10^11^ to 10^12^ RNA copies per g cardiac tissue) compared to HCV [38]. We found a similar pattern in our data with HCV producing the least amount of virus at its peak (~1 PFU/mL/cell), followed by DENV (~10 PFU/mL/cell) and CVB3 (~200 PFU/mL/cell). Considering the RNA synthesis rates, CVB3 is replicating 50-times faster compared to HCV and DENV.

### Broad-spectrum antivirals?

DAAs are highly specific drugs usually designed to inhibit the function of one specific viral protein. Developing broad-spectrum antiviral drugs is challenging. Nevertheless, we were interested in the possibility of a pan-viral drug treatment option. We therefore studied the core processes in the life cycles of our three representatives of plus-strand RNA viruses and administered *in-silico* drugs in mono or combination therapy, with the aim to identify single drug targets or combinations of drug targets that yield an efficient inhibition of all three viruses.

#### Direct acting antivirals against HCV

Several DAAs have been developed and approved for HCV and are able to cure chronic hepatitis C in the majority of patients [120]. DAAs are developed to target one specific protein such as HCV NS3/4A (e.g., first-generation telaprevir or boceprevir and second-/third generation glecaprevir, voxilaprevir and grazoprevir), HCV NS5A (e.g., daclatasvir, velpatasvir, ledipasvir), and HCV NS5B (e.g., sofosbuvir and dasabuvir) [121]. Therefore, the DAAs’ modes of action and efficacies may be used here to validate the results of our *in-silico* drug intervention study. While DAAs blocking HCV NS3/4A intervene with the polyprotein cleavage, HCV NS5A and HCV NS5B inhibitors target the RO formation and vRNA synthesis, respectively [9,59,122]. Our sensitivity and *in-silico* drug analysis suggested high sensitivities for processes associated with HCV RNA replication, which led to an efficient viral reduction by more than 99% with a more than 90% inhibition of the vRNA synthesis rate. Furthermore, our *in-silico* drug analysis predicted that complete HCV NS3/4A inhibition (more than 99.5% polyprotein cleavage inhibition) was necessary to clear the viral load, while in combination with inhibiting vRNA synthesis a combinatory inhibition of more than 90% led to HCV clearance, where viral clearance was mainly driven by inhibiting vRNA synthesis. Our results are in line with current HCV treatment recommendations focusing mainly on a regimen based on a combination of targeting vRNA synthesis alone by inhibiting HCV NS5A and/or NS5B or in combination with HCV NS3/4A, e.g., the combinations of elbasvir (NS5A inhibitor) and grazoprevir (NS3/4A inhibitor), glecaprevir (NS3/4A inhibitor) and pibrentasvir (NS5A inhibitor) or sofosbuvir (NS5B inhibitor) plus velpatasvir (NS5A inhibitor) with the inhibition of NS5A as the backbone of an efficient HCV treatment regimen [123]. Interestingly, the combinatory inhibition of vRNA synthesis and polyprotein cleavage showed pan-viral clearance with the lowest critical efficacies of 0.98, i.e., a 98% inhibition of both processes.

#### Broad-spectrum antivirals and host-directed therapy

The cure of a chronic hepatitis C infection represents a success story for DAAs. However, a subset of HCV patients report treatment failure, severe side effects that impede treatment success, or drug resistance [124]. Targeting cellular components that are crucial for successful and efficient viral replication (so-called host dependency factors) may offer a potential treatment option with a high barrier of resistance. Additionally, plus-strand RNA viruses still represent a major health concern infecting millions of people worldwide, including the viruses in this current study – HCV, DENV and CVB3 – and other plus-strand RNA viruses such as chikungunya, Zika, West Nile, Yellow fever, hepatitis A virus as well as the current global pandemic causing SARS-CoV-2. Even though the identification of pan-serotype antiviral agents is challenging, a DENV inhibitor has been identified, which has shown high efficacy and pan-serotype activity against all known DENV genotypes and serotypes [125]. Our model may serve as a basis towards the development of further virus-specific models as well as pan-viral broad-spectrum antiviral treatment strategies.

Our sensitivity and drug analysis showed that inhibiting translation complex formation, vRNA translation or polyprotein cleavage in combination with vRNA synthesis represent the most promising pan-viral drug targets. As in the case of HCV, targeting vRNA replication and polyprotein cleavage has been highly successful, however, directly targeting the HCV RNA translation (e.g., the HCV IRES RNA structure) or its complex formation is mainly experimental. Another treatment strategy may be targeting host factors hijacked by the virus and involved in almost every process of the viral life cycle [126]. We found that a limitation in the number of available ribosomes may be a key feature limiting efficient virus production due to suppressed host mRNA translation or complete host cell translation shut-off. However, targeting and thus inhibiting the biological function of ribosomes will obviously be challenging and not beneficial for the host. Nevertheless, two proteins were found interacting with vRNA translation: RACK1 and RPS25. Both proteins may be hijacked by DENV and promote DENV mediated cap-independent RNA translation [127]. Additionally, in HCV RACK1 has been shown to inhibit IRES mediated viral RNA translation and viral replication; in the latter case RACK1 binds to HCV NS5A, which induces the formation of ROs [128,129]. Similar to HCV, CVB3 RNA translation is mediated through an IRES and thus RACK1 may be a potential drug target. Furthermore, studying interactions of SARS-CoV-2 proteins with host mRNA identified RACK1 as a binding partner and thus may represent a pan-viral host dependency factor [130].

Interestingly, the very early processes in the viral life cycle, virus entry as well as fusion and release of the vRNA genome, showed significant sensitivities in DENV and CVB3 but was rather negligible in HCV. Further, the release of the viral RNA genome from endosomes showed a higher significant sensitivity compared to viral entry and internalization. Interestingly, cyclophilin A seems to be a host factor involved in the enterovirus A71 (family *Picornaviridae)* fusion/uncoating process and thus vRNA release [131,132]. Furthermore, cyclophilin A inhibitors successfully block or decrease viral replication in a number of plus-strand RNA viruses such as HCV, DENV, West-Nile virus, yellow fever virus, enteroviral A71 and coronavirus [133,134]. Considering that it is involved in both processes that showed highest sensitivities, cyclophilin A may represent a promising pan-viral target [134].

The formation of the replicase complexes represented another sensitive pan-viral process. Replicase complexes are associated with membranes of the ROs either within or outside the RO facing the cytosol [135]. Several studies have shown the significance of host factors in the RO formation being associated with cell permissiveness and vRNA replication efficiency [17,89,118,126]. For example, Tabata et al. (2021) have shown that the RO biogenesis in HCV and SARS-CoV-2 critically depends on the lipid phosphatidic acid synthesis, since inhibiting associated pathways led to an impaired HCV and SARS-CoV-2 RNA replication [136]. However, even though successful in clearing HCV and DENV, in an established infection of a fast-replicating virus such as CVB3, the formation of replicase complexes may not represent an efficient drug target. In steady state, CVB3 replicase complexes are already formed, and the virus cannot be cleared even with a 100% inhibition given for 5 days. Similar results have been found by targeting host factors involved in the formation of replicase complexes of other picornaviruses. Two tested compounds targeting RO formation were not able to block viral replication suggesting that if ROs are already formed, the viral replication continues [137]. Furthermore, targeting host factors involved in RO formation showed lethal cytotoxicity as in the case of PI4KIIIβ and HCV [138]. Interestingly, inhibiting the host factor PI4KB showed that CVB3 RO formation was delayed and CVB3 RNA replication occurred at the Golgi apparatus [139].

Interestingly, incomplete inhibition of some processes may promote viral growth. Our model predicted that targeting viral export from the RO into the cytoplasm in the DENV life cycle led to a 6% increase in virus. Therefore, low-efficacy drugs may lead to the opposite of the desired outcome. Thus, host directed therapy may have a huge potential on the one hand but may result in substantial side effects on the other hand. The identification of host factors with pan-viral activity without lethal toxicity represents a challenge for future research.

### Limitations and outlook

In the current study, we developed the first mathematical model for the intracellular replication of a group of related plus-strand RNA viruses. Even though our model allowed a high degree of parameter identifiability, fit the *in vitro* kinetic data, and is consistent with the current biological knowledge of our studied viruses, there are some weaknesses to consider.

First, our model focuses on a single cell, and hence does not include viral spread. Especially in acute infections with rapidly replicating viruses, viral transmission within organs may be highly relevant to consider. However, since our model was developed for a single step growth curve, we neglected viral spread and focused mainly on intracellular replication processes. Virus-specific mechanisms of viral spread from infected to susceptible cells may be interesting to study in the future.

Second, our experiments were performed in the immuno-compromised Huh7 cell line, we did not consider an intrinsic immune response here. In the future, considering an intrinsic immune response may be an important addition.

Third, even though plus-strand RNA viruses share remarkable similarities in their replication strategy, our model does not consider viruses with more than one open reading frame and ribosomal frameshift. The difference between viruses with one and more open reading frames is the presence of sub-genomic RNA, as in the case of coronaviruses. However, the life cycle of coronaviruses, and in particular SARS-CoV-2, differs from our model by producing non-structural proteins first, followed by viral RNA and sub-genomic RNA synthesis [140]. The sub-genomic RNA is later translated into structural proteins. However, since the core processes of viral non-structural protein production (necessary for vRNA synthesis) and vRNA synthesis itself are common, we do not think that the presence of sub-genomic RNA would have a huge impact on our presented results. Adaptation of the model to coronaviruses is an ongoing topic being followed up in our group.

Fourth, *in vitro* experiments are not a reliable system for an *in vivo* application. Especially our drug treatment study needs experimental validation. However, our model and *in silico* drug analysis showed a high degree of similarity with knowledge and efficacy of DAAs available for HCV.

Fifth, our model has been developed for a one step growth experiment and consequently a single cycle of virus growth. Thus, our model predictions are of a short-term nature and do not study long-term effects.

In summary, in the present study we measured the *in vitro* kinetics of three representatives of plus-strand RNA viruses: HCV, DENV, and CVB3. Based on these experimental measurements, we developed a mathematical model of the intracellular plus-strand RNA virus life cycle. In order to study pan-viral similarities and virus-specific differences, the model was fit simultaneously to the *in vitro* measurements, where the best-fit model was selected based on the AIC and model parameter identifiability. According to our model, the viral life cycles of our three plus-strand RNA representatives differ mainly in processes of viral entry and genome release, the availability of ribosomes involved in viral RNA translation, formation of the replicase complex, and viral trafficking of newly produced viral RNA. Furthermore, our model predicted that the availability of ribosomes involved in viral RNA translation and thus the degree of the host cell translation shut-off may play a key role in acute infection outcome. Interestingly, our modelling predicted that increasing the number of ribosomes available for HCV RNA translation remarkably enhanced the HCV RNA replication efficiency and increased the HCV viral load by three orders of magnitude, a feature we were not able to achieve by increasing the HCV RNA synthesis rate. Furthermore, according to our *in-silico* drug analysis, we found that targeting processes associated with vRNA translation especially polyprotein cleavage together with viral RNA replication substantially decreased viral load and may represent promising drug targets with broad-spectrum antiviral activity.

## Abbreviations

CVB3: Coxsackievirus B3
DAA: Direct acting antivirals
DENV: Dengue virus
d pi: days post infection
HCV: Hepatitis C virus
h pi: hours post infection
ODE: Ordinary differential equations
PV: Poliovirus
RdRp: RNA-dependent RNA polymerase
RO: Replication organelle
ZIKV: Zika virus

## S1 Supporting material: Model selection process

### S1 Supporting data

**S1 Table:** S1 Table: Critical drug efficacy constants in mono and combination therapy and an in-silico drug administration in steady state (100 h pi). For simplicity, we assume that in combination therapy, both drugs have the same efficacy. The lowest critical drug efficacies to clear the virus-specific infection is highlighted in red (TC = translation complex, RC = replicase complex)

| Drug A | Drug B | HCV | DENV | CVB3 |
| --- | --- | --- | --- | --- |
| **TC formation ($k_1$)** | - | 0.96 | 1 | 1 |
| **Translation ($k_2$)** | - | 0.99 | 0.99 | 1 |
| **Polyprotein cleavage ($k_c$)** | - | 0.995 | 1 | 1 |
| **RC formation ($k_{pin}$)** | - | 0.99 | 1 | - |
| **RNA synthesis ($k_{4p}$ and $k_{4m}$)** | - | 0.89 | 0.865 | 0.995 |
| **Viral export ($k_{pout}$)** | - | 1 | 1 | 1 |
| **Virus assembly and release ($k_p$)** | - | 1 | 1 | 1 |
| **TC formation ($k_1$)** | **RNA synthesis ($k_{4p}$ and $k_{4m}$)** | **0.76** | **0.85** | 0.993 |
| **TC formation ($k_1$)** | **RC formation ($k_{pin}$)** | 0.85 | 0.99 | 1 |
| **Translation ($k_2$)** | **RNA synthesis ($k_{4p}$ and $k_{4m}$)** | 0.90 | **0.85** | 0.99 |
| **Translation ($k_2$)** | **RC formation ($k_{pin}$)** | 0.96 | 0.98 | 0.991 |
| **Polyprotein cleavage ($k_c$)** | **RNA synthesis ($k_{4p}$ and $k_{4m}$)** | 0.90 | 0.87 | **0.98** |
| **Polyprotein cleavage ($k_c$)** | **RC formation ($k_{pin}$)** | 0.997 | 0.999 | 1 |

**S1 Figure:**
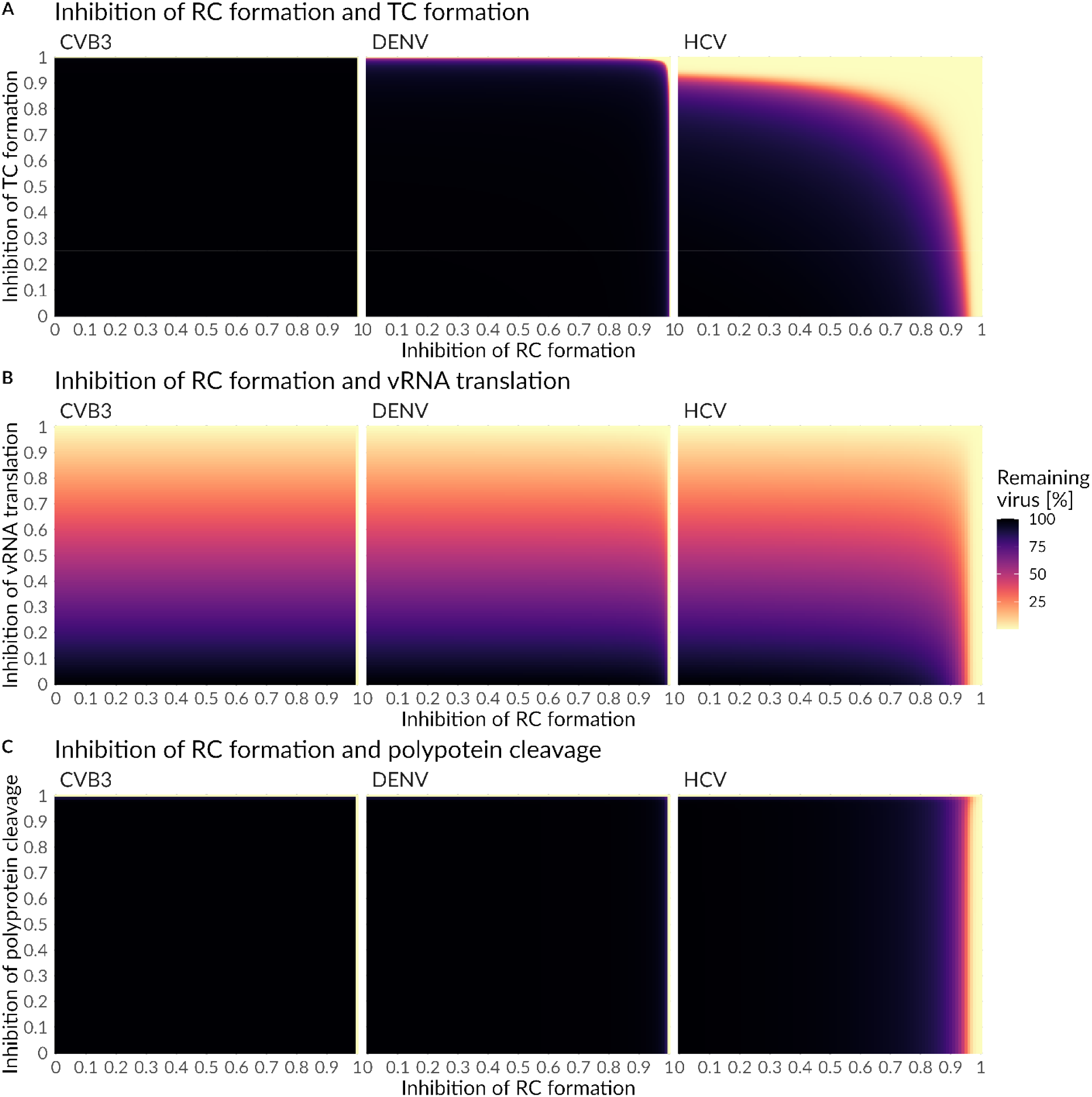
Combined drug effect on **A)** replicase complex (RC) formation and formation of translation complex (TC) **B)** replicase complex (RC) formation and polyprotein cleavage and **C)** replicase complex (RC) formation and vRNA translation and drug administration in steady state (100 h pi). A successful drug treatment leads to a more than 99% viral eradication (light yellow), while an ineffective drug treatment leads to 100% remaining virus (black).

**S2 Figure:**
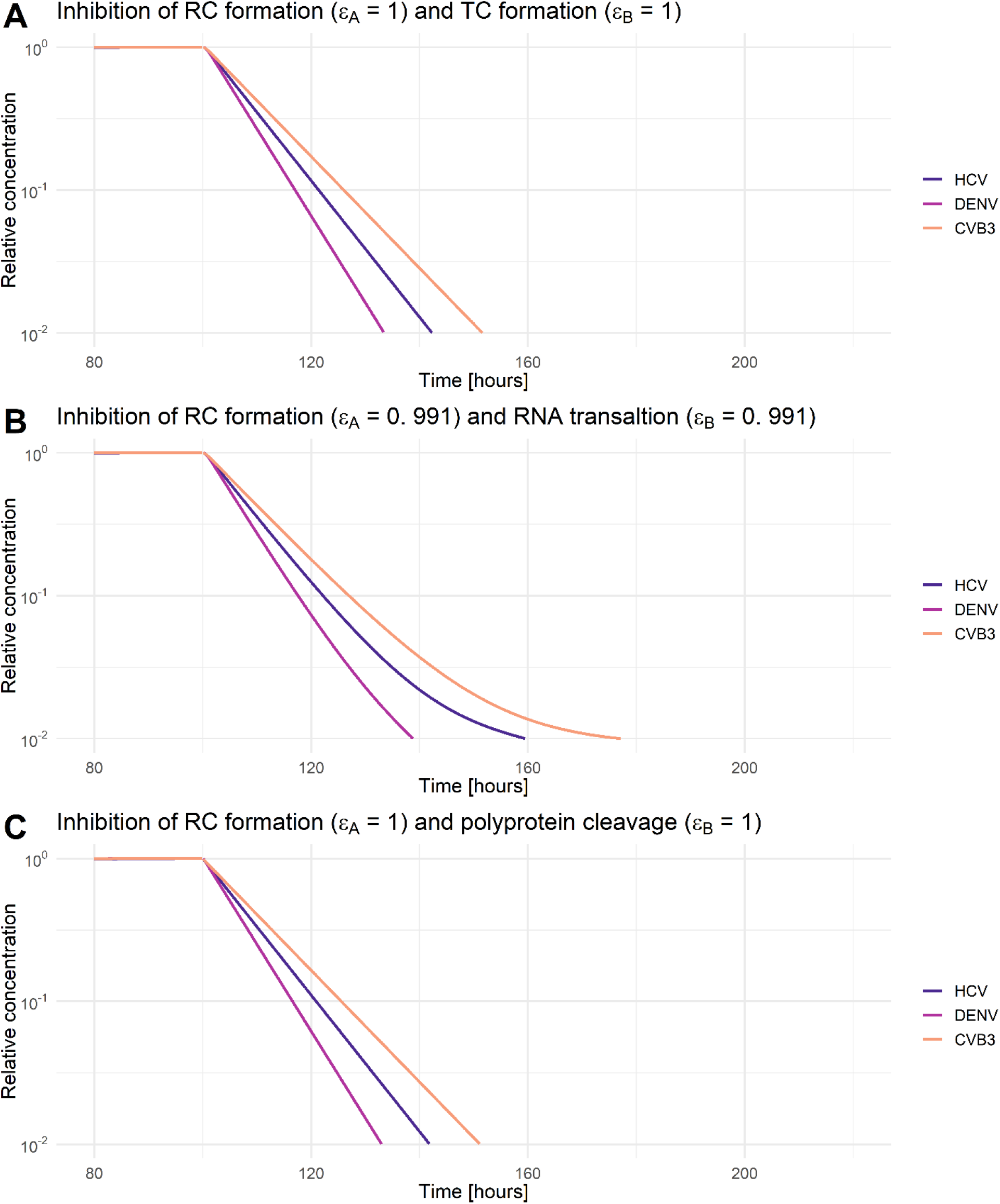
Relative virus decay under combination therapy that clears HCV, DENV, and CVB3 infections. A combined drug effect on **A)** formation of replicase complex (RC) and formation of translation complex (TC), **B)** formation of replicase complex (RC) and translation, and **C)** formation of replicase complex (RC) and polyprotein cleavage. Initiation of treatment was in steady state (100 h pi). The drug efficacy constant (ε_A_ and ε_B_) were chosen as minimal efficacies to clear all three viruses. For comparability, virus-specific concentrations in steady state have been normalized to their virus-specific pre-treatment steady state concentration. A successful drug treatment leads to a more than 99% viral eradication (light yellow), while an ineffective drug treatment leads to 100% remaining virus (black) (see S1 Supporting data).

## Notes

Funding information, This work received funding from the BMBF through the ERASysAPP project SysVirDrug (grant 031A602A). LK received funding from the DFG (grant number KA 2989/13-1). C.D. was supported by a stipend of the DKFZ International PhD Program. Portions of this work were done under the auspices of the U.S. Department of Energy under contract 89233218CNA000001 and supported by NIH grants R01-OD011095, R01-AI078881, and R01-AI116868 to ASP. The funders had no role in study design, data collection and analysis, decision to publish, or preparation of the manuscript.

### Competing Interest Statement

The authors have declared no competing interest.

## References

1. Ciotti M, Angeletti S, Minieri M, Giovannetti M, Benvenuto D, Pascarella S, et al. COVID-19 outbreak: An overview. Chemotherapy. 2020;64: 215–223. doi:10.1159/000507423

2. World Health Organization. WHO coronavirus (COVID-19) dashboard with vaccination data. In: WHO [Internet]. 2021 [cited 7 Mar 2022] pp. 1–5. Available: https://covid19.who.int/

3. Cutler DM, Summers LH. The COVID-19 pandemic and the $16 trillion virus. JAMA – Journal of the American Medical Association. American Medical Association; 2020. pp. 1495–1496. doi:10.1001/jama.2020.19759

4. Shepard DS, Undurraga EA, Halasa YA, Stanaway JD. The global economic burden of dengue: a systematic analysis. Lancet Infect Dis. 2016;16: 935–941. doi:10.1016/S1473-3099(16)00146-8

5. United Nations. A socio-economic impact assessment of the Zika virus in Latin America and the Caribbean. 2017. Available: http://www.undp.org/content/undp/en/home/librarypage/hiv-aids/a-socio-economic-impact-assessment-of-the-zika-virus-in-latin-am.html

6. Barber MJ, Gotham D, Khwairakpam G, Hill A. Price of a hepatitis C cure: Cost of production and current prices for direct-acting antivirals in 50 countries. J Virus Erad. 2020;6: 100001. doi:10.1016/J.JVE.2020.06.001

7. Shakeri A, Srimurugathasan N, Suda KJ, Gomes T, Tadrous M. Spending on Hepatitis C Antivirals in the United States and Canada, 2014 to 2018. Value Heal. 2020;23: 1137–1141. doi:10.1016/J.JVAL.2020.03.021

8. FDA. Coronavirus (COVID-19) update: FDA authorizes first oral antiviral for treatment of COVID-19. In: Food and Drug Administration [Internet]. 2021 p. 1. Available: https://www.fda.gov/news-events/press-announcements/coronavirus-covid-19-update-fda-authorizes-first-oral-antiviral-treatment-covid-19

9. Hayes CN, Imamura M, Tanaka J, Chayama K. Road to elimination of HCV: Clinical challenges in HCV management. Liver International. John Wiley & Sons, Ltd; 2022. doi:10.1111/liv.15150

10. World Health Organization. Hepatitis C. 2019. Available: https://www.who.int/news-room/fact-sheets/detail/hepatitis-c

11. World Health Organization. Dengue and severe dengue. 2016. doi:10.1111/1469-0691.12442

12. Colpitts CC, El-Saghire H, Pochet N, Schuster C, Baumert TF. High-throughput approaches to unravel hepatitis C virus-host interactions. Virus Res. 2016;218: 18–24. doi:10.1016/j.virusres.2015.09.013

13. Genoni A, Canducci F, Rossi A, Broccolo F, Chumakov K, Bono G, et al. Revealing enterovirus infection in chronic human disorders: An integrated diagnostic approach. Sci Rep. 2017;7: 5013. doi:10.1038/s41598-017-04993-y

14. Baggen J, Thibaut HJ, Strating JRPM, Van Kuppeveld FJM. The life cycle of non-polio enteroviruses and how to target it. Nature Reviews Microbiology. Nat Rev Microbiol; 2018. pp. 368–381. doi:10.1038/s41579-018-0005-4

15. Garmaroudi FS, Marchant D, Hendry R, Luo H, Yang D, Ye X, et al. Coxsackievirus B3 replication and pathogenesis. 2015;10: 629–652.

16. Romero-Brey I, Merz A, Chiramel A, Lee JY, Chlanda P, Haselman U, et al. Three-dimensional architecture and biogenesis of membrane structures associated with hepatitis C virus replication. Luo GG, editor. PLoS Pathog. 2012;8: e1003056. doi:10.1371/journal.ppat.1003056

17. Belov GA, van Kuppeveld FJ. (+)RNA viruses rewire cellular pathways to build replication organelles. Curr Opin Virol. 2012;2: 740–747. doi:10.1016/j.coviro.2012.09.006

18. Miller S, Krijnse-Locker J. Modification of intracellular membrane structures for virus replication. Nat Rev Microbiol. 2008;6: 363–374. doi:10.1038/nrmicro1890

19. Binder M, Sulaimanov N, Clausznitzer D, Schulze M, Hüber CM, Lenz SM, et al. Replication vesicles are load-and choke-points in the hepatitis C virus lifecycle. PLoS Pathog. 2013;9: e1003561. doi:10.1371/journal.ppat.1003561

20. Paul D, Bartenschlager R. Architecture and biogenesis of plus-strand RNA virus replication factories. World J Virol. 2013;2: 32–48. doi:10.5501/wjv.v2.i2.32

21. Limpens RWAL, van der Schaar HM, Kumar D, Koster AJ, Snijder EJ, van Kuppeveld FJM, et al. The transformation of enterovirus replication structures: A three-dimensional study of single-and double-membrane compartments. MBio. 2011;2. doi:10.1128/mBio.00166-11

22. Gale M, Tan SL, Katze MG. Translational control of viral gene expression in eukaryotes. Microbiol Mol Biol Rev. 2000;64: 239–80. doi:10.1128/MMBR.64.2.239-280.2000

23. Roth H, Magg V, Uch F, Mutz P, Klein P, Haneke K, et al. Flavivirus infection uncouples translation suppression from cellular stress responses. MBio. 2017;8. doi:10.1128/mBio.02150-16

24. Huang J-Y, Su W-C, Jeng K-S, Chang T-H, Lai MMC. Attenuation of 40S ribosomal subunit abundance differentially affects host and HCV translation and suppresses HCV replication. PLoS Pathog. 2012;8: e1002766. doi:10.1371/journal.ppat.1002766

25. Zitzmann C, Kaderali L. Mathematical analysis of viral replication dynamics and antiviral treatment strategies: From basic models to age-based multi-scale modeling. Front Microbiol. Frontiers; 2018. p. 1546. doi:10.3389/fmicb.2018.01546

26. Perelson AS, Ke R. Mechanistic modelling of SARS-CoV-2 and other infectious diseases and the effects of therapeutics. Clin Pharmacol Ther. 2021. doi:10.1002/cpt.2160

27. Layden TJ, Layden JE, Ribeiro RM, Perelson AS. Mathematical modeling of viral kinetics: A tool to understand and optimize therapy. Clin Liver Dis. 2003;7: 163–178. doi:10.1016/S1089-3261(02)00063-6

28. Perelson AS, Ribeiro RM. Hepatitis B virus kinetics and mathematical modeling. Semin Liver Dis. 2004;24: 11–16. doi:10.1055/s-2004-828673

29. Smith AM, Perelson AS. Influenza A virus infection kinetics: Quantitative data and models. Wiley Interdiscip Rev Syst Biol Med. 2011;3: 429–445. doi:10.1002/wsbm.129

30. Bonhoeffer S, Coffin JM, Nowak MA. Human Immunodeficiency Virus Drug Therapy and Virus Load. J Virol. 1997;71: 3275–3278.

31. Perelson AS, Ribeiro RM. Modeling the within-host dynamics of HIV infection. BMC Biol. 2013;11: 96. doi:10.1186/1741-7007-11-96

32. Tuiskunen Bäck A, Lundkvist Å. Dengue viruses–an overview. Infect Ecol Epidemiol. 2013;3: 19839. doi:10.3402/iee.v3i0.19839

33. Moradpour D, Penin F, Rice CM. Replication of hepatitis C virus. J Gen Virol. 2007;5: 453–463. doi:10.1038/nrmicro1645

34. Chen BS, Lee HC, Lee KM, Gong YN, Shih SR. Enterovirus and encephalitis. Frontiers in Microbiology. Frontiers Media S.A.; 2020. p. 261. doi:10.3389/fmicb.2020.00261

35. Koestner W, Spanier J, Klause T, Tegtmeyer P-K, Becker J, Herder V, et al. Interferon-beta expression and type I interferon receptor signaling of hepatocytes prevent hepatic necrosis and virus dissemination in Coxsackievirus B3-infected mice. Lemon SM, editor. PLOS Pathog. 2018;14: e1007235. doi:10.1371/journal.ppat.1007235

36. WHO. Hepatitis C. 2019 [cited 20 Aug 2019]. Available: https://www.who.int/news-room/fact-sheets/detail/hepatitis-c

37. J. E Cogan. Dengue and severe dengue. In: World Health Organization [Internet]. 2018. Available: https://www.who.int/news-room/fact-sheets/detail/dengue-and-severe-dengue

38. Reetoo KN, Osman SA, Illavia SJ, Cameron-Wilson CL, Banatvala JE, Muir P. Quantitative analysis of viral RNA kinetics in coxsackievirus B3-induced murine myocarditis: Biphasic pattern of clearance following acute infection, with persistence of residual viral RNA throughout and beyond the inflammatory phase of disease. J Gen Virol. 2000;81: 2755–2762. doi:10.1099/0022-1317-81-11-2755

39. Pybus OG, Charleston MA, Gupta S, Rambaut A, Holmes EC, Harvey PH. The epidemic behavior of the hepatitis C virus. Science (80-). 2001;292: 2323–2325. doi:10.1126/science.1058321

40. Liu Y, Lillepold K, Semenza JC, Tozan Y, Quam MBM, Rocklöv J. Reviewing estimates of the basic reproduction number for dengue, Zika and chikungunya across global climate zones. Environmental Research. Academic Press Inc.; 2020. p. 109114. doi:10.1016/j.envres.2020.109114

41. Lim CTK, Jiang L, Ma S, James L, Ang LW. Basic reproduction number of coxsackievirus type A6 and A16 and enterovirus 71: Estimates from outbreaks of hand, foot and mouth disease in Singapore, a tropical city-state. Epidemiol Infect. 2016;144: 1028–1034. doi:10.1017/S0950268815002137

42. Ma E, Fung C, Yip SHL, Wong C, Chuang SK, Tsang T. Estimation of the basic reproduction number of enterovirus 71 and coxsackievirus A16 in hand, foot, and mouth disease outbreaks. Pediatr Infect Dis J. 2011;30: 675–679. doi:10.1097/INF.0b013e3182116e95

43. Martinelli A de LC, Brown D, Morris A, Dhillon A, Dayley P, Dusheiko G. Quantitation of HCV RNA in liver of patients with chronic hepatitis C. Arq Gastroenterol. 2000;37: 203–207. doi:10.1590/S0004-28032000000400003

44. Ben-Shachar R, Koelle K. Transmission-clearance trade-offs indicate that dengue virulence evolution depends on epidemiological context. Nat Commun. 2018;9: 2355. doi:10.1038/s41467-018-04595-w

45. Major ME, Dahari H, Mihalik K, Puig M, Rice CM, Neumann AU, et al. Hepatitis C virus kinetics and host responses associated with disease and outcome of infection in chimpanzees. Hepatology. 2004;39: 1709–1720. doi:10.1002/hep.20239

46. Nainan O V., Alter MJ, Kruszon-Moran D, Gao FX, Xia G, McQuillan G, et al. Hepatitis C virus genotypes and viral concentrations in participants of a general population survey in the United States. Gastroenterology. 2006;131: 478–484. doi:10.1053/j.gastro.2006.06.007

47. Hajarizadeh B, Grady B, Page K, Kim AY, McGovern BH, Cox AL, et al. Patterns of hepatitis C Virus RNA levels during acute infection: The InC3 study. Blackard J, editor. PLoS One. 2015;10: e0122232. doi:10.1371/journal.pone.0122232

48. Cherry JD, Krogstad P. Enterovirus and parechovirus infections. Infectious Diseases of the Fetus and Newborn Infant. W.B. Saunders; 2011. pp. 756–799. doi:10.1016/B978-1-4160-6400-8.00024-9

49. Koutsoudakis G, Herrmann E, Kallis S, Bartenschlager R, Pietschmann T. The level of CD81 cell surface expression is a key determinant for productive entry of hepatitis C virus into host cells. J Virol. 2007;81: 588–598. doi:10.1128/jvi.01534-06

50. Dächert C, Gladilin E, Binder M. Gene expression profiling of different HuH7 variants reveals novel hepatitis C virus host factors. Viruses. 2019;12. doi:10.3390/v12010036

51. P C, N S. Single-step method of RNA isolation by acid guanidinium thiocyanate-phenol-chloroform extraction. Anal Biochem. 1987;162: 156–159. doi:10.1006/ABIO.1987.9999

52. Grünvogel O, Colasanti O, Lee JY, Klöss V, Belouzard S, Reustle A, et al. Secretion of hepatitis C virus replication intermediates reduces activation of toll-like receptor 3 in hepatocytes. Gastroenterology. 2018;154: 2237–2251.e16. doi:10.1053/j.gastro.2018.03.020

53. Lanke KHW, van der Schaar HM, Belov GA, Feng Q, Duijsings D, Jackson CL, et al. GBF1, a guanine nucleotide exchange factor for Arf, is crucial for coxsackievirus B3 RNA replication. J Virol. 2009;83: 11940–9. doi:10.1128/JVI.01244-09

54. Feng Q, Hato SV., Langereis MA, Zoll J, Virgen-Slane R, Peisley A, et al. MDA5 detects the double-stranded RNA replicative form in picornavirus-infected cells. Cell Rep. 2012;2: 1187–1196. doi:10.1016/j.celrep.2012.10.005

55. Zitzmann C, Schmid B, Ruggieri A, Perelson AS, Binder M, Bartenschlager R, et al. A coupled mathematical model of the intracellular replication of dengue virus and the host cell immune response to infection. Front Microbiol. 2020;11: 725. doi:10.3389/fmicb.2020.00725

56. Schmid B, Rinas M, Ruggieri A, Acosta EG, Bartenschlager M, Reuter A, et al. Live cell analysis and mathematical modeling identify determinants of attenuation of dengue virus 2-O-methylation mutant. PLoS Pathog. 2015;11: e1005345. doi:10.1371/journal.ppat.1005345

57. Zitzmann C, Kaderali L, Perelson AS. Mathematical modeling of hepatitis C RNA replication, exosome secretion and virus release. PLoS Comput Biol. 2020;16: e1008421. doi:10.1371/journal.pcbi.1008421

58. Kazakov T, Yang F, Ramanathan HN, Kohlway A, Diamond MS, Lindenbach BD. Hepatitis C virus RNA replication depends on specific cis-and trans-acting activities of viral nonstructural proteins. PLoS Pathog. 2015;11: e1004817. doi:10.1371/journal.ppat.1004817

59. Benzine T, Brandt R, Lovell WC, Yamane D, Neddermann P, De Francesco R, et al. NS5A inhibitors unmask differences in functional replicase complex half-life between different hepatitis C virus strains. Randall G, editor. PLOS Pathog. 2017;13: e1006343. doi:10.1371/journal.ppat.1006343

60. Heldt FS, Frensing T, Reichl U. Modeling the intracellular dynamics of influenza virus replication to understand the control of viral RNA synthesis. J Virol. 2012;86: 7806–17. doi:10.1128/JVI.00080-12

61. Laske T, Heldt FS, Hoffmann H, Frensing T, Reichl U. Modeling the intracellular replication of influenza A virus in the presence of defective interfering RNAs. Virus Res. 2016;213: 90–99. doi:10.1016/j.virusres.2015.11.016

62. Raue A, Steiert B, Schelker M, Kreutz C, Maiwald T, Hass H, et al. Data2Dynamics: a modeling environment tailored to parameter estimation in dynamical systems. Bioinformatics. 2015;31: 3558–3560. doi:10.1093/bioinformatics/btv405

63. Raue A, Kreutz C, Maiwald T, Bachmann J, Schilling M, Klingmüller U, et al. Structural and practical identifiability analysis of partially observed dynamical models by exploiting the profile likelihood. Bioinformatics. 2009;25: 1923–1929. doi:10.1093/bioinformatics/btp358

64. Marino S, Hogue IB, Ray CJ, Kirschner DE. A methodology for performing global uncertainty and sensitivity analysis in systems biology. J Theor Biol. 2008;254: 178–96. doi:10.1016/j.jtbi.2008.04.011

65. Aunins TR, Marsh KA, Subramanya G, Uprichard SL, Perelson AS, Chatterjee A. Intracellular hepatitis C modeling predicts infection dynamics and viral protein mechanisms. J Virol. 2018;92: JVI.02098–17. doi:10.1128/JVI.02098-17

66. Regoes RR, Crotty S, Antia R, Tanaka MM. Optimal replication of poliovirus within cells. Am Nat. 2005;165: 364–73. doi:10.1086/428295

67. Byk LA, Iglesias NG, De Maio FA, Gebhard LG, Rossi M, Gamarnik A V. Dengue virus genome uncoating requires ubiquitination. MBio. 2016;7: e00804–16. doi:10.1128/mBio.00804-16

68. Simoes EA, Sarnow P. An RNA hairpin at the extreme 5’ end of the poliovirus RNA genome modulates viral translation in human cells. J Virol. 1991;65: 913–921. doi:10.1128/jvi.65.2.913-921.1991

69. Gohara DW, Arnold JJ, Cameron CE. Poliovirus RNA-dependent RNA polymerase (3Dpol): Kinetic, thermodynamic, and structural analysis of ribonucleotide selection. Biochemistry. 2004;43: 5149–5158. doi:10.1021/bi035429s

70. Goo L, Dowd KA, Smith ARY, Pelc RS, Demaso CR, Pierson TC. Zika virus is not uniquely stable at physiological temperatures compared to other flaviviruses. MBio. 2016;7. doi:10.1128/mBio.01396-16

71. Carson SD, Hafenstein S, Lee H. MOPS and coxsackievirus B3 stability. Virology. 2017;501: 183–187. doi:10.1016/j.virol.2016.12.002

72. Carson SD, Chapman NM, Hafenstein S, Tracy S. Variations of coxsackievirus B3 capsid primary structure, ligands, and stability Are selected for in a coxsackievirus and adenovirus receptor-limited environment. J Virol. 2011;85: 3306–3314. doi:10.1128/jvi.01827-10

73. Persaud M, Martinez-Lopez A, Buffone C, Porcelli SA, Diaz-Griffero F. Infection by Zika viruses requires the transmembrane protein AXL, endocytosis and low pH. Virology. 2018;518: 301–312. doi:10.1016/j.virol.2018.03.009

74. Chatel-Chaix L, Bartenschlager R. Dengue virus-and hepatitis C virus-induced replication. J Virol. 2014;88: 5907–5911.

75. Perelson AS, Neumann AU, Markowitz M, Leonard JM, Ho DD. HIV-1 dynamics in vivo: virion clearance rate, infected cell life-span, and viral generation time. Science (80-). 1996;271: 1582–1586. doi:10.1126/science.271.5255.1582

76. Neumann AU, Lam NP, Dahari H, Gretch DR, Wiley TE, Layden TJ, et al. Hepatitis C viral dynamics in vivo and the antiviral efficacy of interferon-α therapy. Science (80-). 1998;282: 103–107. doi:10.1126/science.282.5386.103

77. Perelson AS, Essunger P, Cao Y, Vesanen M, Hurley A, Saksela K, et al. Decay characteristics of HIV-1-infected compartments during combination therapy. Nature. 1997;387: 188–191. doi:10.1038/387188a0

78. Baccam P, Beauchemin C, Macken CA, Hayden FG, Perelson AS. Kinetics of influenza A virus infection in humans. J Virol. 2006;80: 7590–9. doi:10.1128/JVI.01623-05

79. Dahari H, Ribeiro RM, Rice CM, Perelson AS. Mathematical modeling of subgenomic hepatitis C virus replication in Huh-7 cells. J Virol. 2007;81: 750–60. doi:10.1128/JVI.01304-06

80. Quintela B de M, Conway JM, Hyman JM, Guedj J, dos Santos RW, Lobosco M, et al. A new age-structured multiscale model of the hepatitis C virus life-cycle during infection and therapy with direct-acting antiviral agents. Front Microbiol. 2018;9: 601. doi:10.3389/fmicb.2018.00601

81. Reddy B, Yin J. Quantitative intracellular kinetics of HIV type 1. AIDS Res Hum Retroviruses. 1999;15: 273–283. doi:10.1089/088922299311457

82. Heldt FS, Frensing T, Pflugmacher A, Gröpler R, Peschel B, Reichl U. Multiscale modeling of influenza A virus infection supports the development of direct-acting antivirals. Koelle K, editor. PLoS Comput Biol. 2013;9: e1003372. doi:10.1371/journal.pcbi.1003372

83. Reichl U, Sidorenko Y. Dynamics of virus-host cell interaction. Bioinformatics-From Genomes to Therapies. Weinheim, Germany: Wiley-VCH Verlag GmbH; 2008. pp. 861–898. doi:10.1002/9783527619368.ch23

84. Frensing T, Heldt FS, Pflugmacher A, Behrendt I, Jordan I, Flockerzi D, et al. Continuous influenza virus production in cell culture shows a periodic accumulation of defective interfering particles. Pöhlmann S, editor. PLoS One. 2013;8: e72288. doi:10.1371/journal.pone.0072288

85. Heldt FS, Kupke SY, Dorl S, Reichl U, Frensing T. Single-cell analysis and stochastic modelling unveil large cell-to-cell variability in influenza A virus infection. Nat Commun. 2015;6: 8938. doi:10.1038/ncomms9938

86. Sidorenko Y, Voigt A, Schulze-Horsel J, Reichl U, Kienle A. Stochastic population balance modeling of influenza virus replication in vaccine production processes. II. Detailed description of the replication mechanism. Chem Eng Sci. 2008. doi:10.1016/j.ces.2007.12.034

87. Sidorenko Y, Reichl U. Structured model of influenza virus replication in MDCK cells. Biotechnol Bioeng. 2004;88: 1–14. doi:10.1002/bit.20096

88. Chhajer H, Rizvi VA, Roy R. Life cycle process dependencies of positive-sense RNA viruses suggest strategies for inhibiting productive cellular infection. J R Soc Interface. 2021;18. doi:10.1098/RSIF.2021.0401

89. Baggen J, Thibaut HJ, Strating JRPM, Van Kuppeveld FJM. The life cycle of non-polio enteroviruses and how to target it. Nature Reviews Microbiology. Nature Publishing Group; 2018. pp. 368–381. doi:10.1038/s41579-018-0005-4

90. Lohmann V, Bartenschlager R. On the history of hepatitis C virus cell culture systems. J Med Chem. 2014;57: 1627–1642. doi:10.1021/JM401401N

91. Fischl W, Bartenschlager R. Exploitation of cellular pathways by Dengue virus. Current Opinion in Microbiology. 2011. pp. 470–475. doi:10.1016/j.mib.2011.07.012

92. Clyde K, Kyle JL, Harris E. Recent advances in deciphering viral and host determinants of dengue virus replication and pathogenesis. J Virol. 2006;80: 11418–11431. doi:10.1128/jvi.01257-06

93. Anderson R. Manipulation of cell surface macromolecules by flaviviruses. Adv Virus Res. 2003;59: 229–274. doi:10.1016/S0065-3527(03)59007-8

94. Hsu NY, Ilnytska O, Belov G, Santiana M, Chen YH, Takvorian PM, et al. Viral reorganization of the secretory pathway generates distinct organelles for RNA replication. Cell. 2010;141: 799–811. doi:10.1016/j.cell.2010.03.050

95. Bushell M, Sarnow P. Hijacking the translation apparatus by RNA viruses. Journal of Cell Biology. The Rockefeller University Press; 2002. pp. 395–399. doi:10.1083/jcb.200205044

96. Summers DF, Maizel J V., Darnell JE. The decrease in size and synthetic activity of poliovirus polysomes late in the infectious cycle. Virology. 1967;31: 427–435. doi:10.1016/0042-6822(67)90222-X

97. Boersma S, Rabouw HH, Bruurs LJM, Pavlovič T, van Vliet ALW, Beumer J, et al. Translation and replication dynamics of single RNA viruses. Cell. 2020;183: 1930–1945.e23. doi:10.1016/j.cell.2020.10.019

98. Reid DW, Campos RK, Child JR, Zheng T, Chan KWK, Bradrick SS, et al. Dengue virus selectively annexes endoplasmic reticulum-associated translation machinery as a strategy for co-opting host cell protein synthesis. J Virol. 2018;92: 1766–1783. doi:10.1128/jvi.01766-17

99. Melia CE, Peddie CJ, de Jong AWM, Snijder EJ, Collinson LM, Koster AJ, et al. Origins of enterovirus replication organelles established by whole-cell electron microscopy. MBio. 2019;10. doi:10.1128/mbio.00951-19

100. Melia CE, van der Schaar HM, Lyoo H, Limpens RWAL, Feng Q, Wahedi M, et al. Escaping host factor PI4KB inhibition: Enterovirus genomic RNA replication in the absence of replication organelles. Cell Rep. 2017;21: 587–599. doi:10.1016/j.celrep.2017.09.068

101. Li X, Wang M, Cheng A, Wen X, Ou X, Mao S, et al. Enterovirus replication o organelles and inhibitors of their formation. Frontiers in Microbiology. Frontiers Media S.A.; 2020. p. 1817. doi:10.3389/fmicb.2020.01817

102. Iglesias NG, Gamarnik A V. RNA Biology Dynamic RNA structures in the dengue virus genome. 2011 [cited 6 Aug 2020]. doi:10.4161/rna.8.2.14992

103. Villordo SM, Alvarez DE, Gamarnik AV. A balance between circular and linear forms of the dengue virus genome is crucial for viral replication. RNA. 2010;16: 2325–2335. doi:10.1261/rna.2120410

104. Bolten R, Egger D, Gosert R, Schaub G, Landmann L, Bienz K. Intracellular localization of poliovirus plus-and minus-Strand RNA visualized by strand-specific fluorescent in situ hybridization. J Virol. 1998;72: 8578–8585. doi:10.1128/jvi.72.11.8578-8585.1998

105. Guo J-T, Bichko V V, Seeger C. Effect of alpha interferon on the hepatitis C virus replicon. J Virol. 2001;75: 8516–8523. doi:10.1128/jvi.75.18.8516-8523.2001

106. Quinkert D, Bartenschlager R, Lohmann V. Quantitative analysis of the hepatitis C virus replication complex. J Virol. 2005;79: 13594–13605. doi:10.1128/jvi.79.21.13594-13605.2005

107. Iwasaki A, Medzhitov R. Innate responses to viral infections. 6th ed. In: Fields BN, Knipe DM, Howley PM, editors. Fields Virology: Sixth Edition. 6th ed. Wolters Kluwer Health/Lippincott Williams & Wilkins; 2013. pp. 189–213.

108. Boersma S, Rabouw HH, Bruurs LJM, Pavlovič T, van Vliet ALW, Beumer J, et al. Translation and replication dynamics of single RNA viruses. Cell. 2020;183: 1930–1945.e23. doi:10.1016/j.cell.2020.10.019

109. Lohmann V, Körner F, Koch JO, Herian U, Theilmann L, Bartenschlager R. Replication of subgenomic hepatitis C virus RNAs in a hepatoma cell line. Science (80-). 1999;285: 110–113. doi:10.1126/science.285.5424.110

110. Oh J-W, Ito T, Lai MMC. A recombinant hepatitis C virus RNA-dependent RNA polymerase capable of copying the full-length viral RNA. J Virol. 1999;73: 7694–7702. doi:10.1128/jvi.73.9.7694-7702.1999

111. Ma H, Leveque V, De Witte A, Li W, Hendricks T, Clausen SM, et al. Inhibition of native hepatitis C virus replicase by nucleotide and non-nucleoside inhibitors. Virology. 2005;332: 8–15. doi:10.1016/j.virol.2004.11.024

112. Tan BH, Fu J, Sugrue RJ, Yap EH, Chan YC, Tan YH. Recombinant dengue type 1 virus NS5 protein expressed in Escherichia coli exhibits RNA-dependent RNA polymerase activity. Virology. 1996;216: 317–325. doi:10.1006/viro.1996.0067

113. Yang Y, Wang Z. IRES-mediated cap-independent translation, a path leading to hidden proteome. J Mol Cell Biol. 2019;11: 911–919. doi:10.1093/JMCB/MJZ091

114. Lee KM, Chen CJ, Shih SR. Regulation mechanisms of viral IRES-driven translation. Trends in Microbiology. Elsevier; 2017. pp. 546–561. doi:10.1016/j.tim.2017.01.010

115. Pelletier J, Sonenberg N. The organizing principles of eukaryotic ribosome recruitment. Annual Review of Biochemistry. 2019. pp. 307–335. doi:10.1146/annurev-biochem-013118-111042

116. Fernández-García L, Angulo J, Ramos H, Barrera A, Pino K, Vera-Otarola J, et al. The internal ribosome entry site of dengue virus mRNA is Active when cap-dependent translation initiation is inhibited. J Virol. 2021;95. doi:10.1128/jvi.01998-20

117. Finkel Y, Gluck A, Nachshon A, Winkler R, Fisher T, Rozman B, et al. SARS-CoV-2 uses a multipronged strategy to impede host protein synthesis. Nature. 2021;594: 240–245. doi:10.1038/s41586-021-03610-3

118. de Wilde AH, Snijder EJ, Kikkert M, van Hemert MJ. Host factors in coronavirus replication. Current Topics in Microbiology and Immunology. Springer Verlag; 2018. pp. 1–42. doi:10.1007/82_2017_25

119. Ribeiro RM, Li H, Wang S, Stoddard MB, Learn GH, Korber BT, et al. Quantifying the diversification of hepatitis C virus (HCV) during primary infection: Estimates of the in vivo mutation rate. PLoS Pathog. 2012;8. doi:10.1371/journal.ppat.1002881

120. Li DK, Chung RT. Overview of direct-acting antiviral drugs and drug resistance of hepatitis C virus. Methods in Molecular Biology. Humana Press Inc.; 2019. pp. 3–32. doi:10.1007/978-1-4939-8976-8_1

121. Perales C, Quer J, Gregori J, Esteban JI, Domingo E. Resistance of hepatitis C virus to inhibitors: Complexity and clinical implications. Viruses. MDPI AG; 2015. pp. 5746–5766. doi:10.3390/v7112902

122. McGivern DR, Masaki T, Williford S, Ingravallo P, Feng Z, Lahser F, et al. Kinetic analyses reveal potent and early blockade of hepatitis C virus assembly by NS5A inhibitors. Gastroenterology. 2014;147. doi:10.1053/J.GASTRO.2014.04.021

123. Bhattacharjee C, Singh M, Das D, Chaudhuri S, Mukhopadhyay A. Current therapeutics against HCV. VirusDisease. 2021;32: 228. doi:10.1007/S13337-021-00697-0

124. Alazard-Dany N, Denolly S, Boson B, Cosset FL. Overview of hcv life cycle with a special focus on current and possible future antiviral targets. Viruses. MDPI AG; 2019. p. 30. doi:10.3390/v11010030

125. Kaptein SJF, Goethals O, Kiemel D, Marchand A, Kesteleyn B, Bonfanti JF, et al. A pan-serotype dengue virus inhibitor targeting the NS3–NS4B interaction. Nat 2021 5987881. 2021;598: 504–509. doi:10.1038/s41586-021-03990-6

126. Nagy PD, Pogany J. The dependence of viral RNA replication on co-opted host factors. Nat Rev Microbiol. 2012;10: 137–149. doi:10.1038/nrmicro2692

127. Hafirassou ML, Meertens L, Umaña-Diaz C, Labeau A, Dejarnac O, Bonnet-Madin L, et al. A global interactome map of the dengue virus NS1 identifies virus restriction and dependency host factors. Cell Rep. 2017;21: 3900–3913. doi:10.1016/j.celrep.2017.11.094

128. Lee JS, Tabata K, Twu WI, Rahman MS, Kim HS, Yu JB, et al. RACK1 mediates rewiring of intracellular networks induced by hepatitis C virus infection. PLoS Pathog. 2019;15: e1008021. doi:10.1371/journal.ppat.1008021

129. Majzoub K, Hafirassou ML, Meignin C, Goto A, Marzi S, Fedorova A, et al. RACK1 controls IRES-mediated translation of viruses. Cell. 2014;159: 1086–1095. doi:10.1016/j.cell.2014.10.041

130. Adams DR, Ron D, Kiely PA. RACK1, A multifaceted scaffolding protein: Structure and function. Cell Commun Signal. 2011;9: 22. doi:10.1186/1478-811X-9-22

131. Kobayashi K, Koike S. Cellular receptors for enterovirus A71. J Biomed Sci 2020 271. 2020;27: 1–12. doi:10.1186/S12929-020-0615-9

132. Qing J, Wang Y, Sun Y, Huang J, Yan W, Wang J, et al. Cyclophilin A associates with enterovirus-71 virus capsid and plays an essential role in viral infection as an uncoating regulator. PLoS Pathog. 2014;10: e1004422. doi:10.1371/journal.ppat.1004422

133. Dawar FU, Tu J, Khattak MNK, Mei J, Lin L. Cyclophilin a: A key factor in virus replication and potential target for anti-viral therapy. Curr Issues Mol Biol. 2017;21: 1–20. doi:10.21775/cimb.021.001

134. Bauer L, Lyoo H, van der Schaar HM, Strating JR, van Kuppeveld FJ. Direct-acting antivirals and host-targeting strategies to combat enterovirus infections. Current Opinion in Virology. Elsevier B.V.; 2017. pp. 1–8. doi:10.1016/j.coviro.2017.03.009

135. Paul D, Hoppe S, Saher G, Krijnse-Locker J, Bartenschlager R. Morphological and biochemical characterization of the membranous hepatitis C virus replication compartment. J Virol. 2013;87: 10612–27. doi:10.1128/JVI.01370-13

136. Tabata K, Prasad V, Paul D, Lee JY, Pham MT, Twu WI, et al. Convergent use of phosphatidic acid for hepatitis C virus and SARS-CoV-2 replication organelle formation. Nat Commun 2021 121. 2021;12: 1–15. doi:10.1038/s41467-021-27511-1

137. Ford Siltz LA, Viktorova EG, Zhang B, Kouiavskaia D, Dragunsky E, Chumakov K, et al. New small-molecule inhibitors effectively blocking picornavirus replication. J Virol. 2014;88: 11091–11107. doi:10.1128/jvi.01877-14

138. LaMarche MJ, Borawski J, Bose A, Capacci-Daniel C, Colvin R, Dennehy M, et al. Anti-hepatitis C virus activity and toxicity of type III phosphatidylinositol-4-kinase beta inhibitors. Antimicrob Agents Chemother. 2012;56: 5149–5156. doi:10.1128/AAC.00946-12

139. Melia CE, van der Schaar HM, Lyoo H, Limpens RWAL, Feng Q, Wahedi M, et al. Escaping host factor PI4KB inhibition: Enterovirus genomic RNA replication in the absence of replication organelles. Cell Rep. 2017;21: 587–599. doi:10.1016/j.celrep.2017.09.068

140. V’kovski P, Kratzel A, Steiner S, Stalder H, Thiel V. Coronavirus biology and replication: implications for SARS-CoV-2. Nature Reviews Microbiology. Nature Publishing Group; 2021. pp. 155–170. doi:10.1038/s41579-020-00468-6

